# The size-weight illusion and beyond: a new model of perceived weight

**DOI:** 10.1101/2024.09.01.610669

**Authors:** Veronica Pisu, Erich W. Graf, Wendy J. Adams

## Abstract

In the size-weight illusion (SWI), the smaller of two same-weight, same apparent material objects is perceived as heavier. The SWI has proved difficult to explain via traditional Bayesian models, which predict the opposite effect: expected weight from size (smaller = lighter) should be integrated with felt weight, such that the smaller object should be perceptually lighter. Other authors have proposed that weight and density are combined according to Bayesian principles, or that Bayesian models incorporating efficient coding can predict the SWI via ‘likelihood repulsion’. These models, however, have been evaluated only under the narrow conditions of typical SWI stimuli. Here we establish a general model of perceived weight for pairs of objects that differ in weight and / or density and / or size by varying amounts. In a visuo-haptic task, participants (N = 30) grasped and lifted pairs of cubes, and reported their perceived heaviness. We report that the SWI occurs even at very small density differences, repudiating the idea that the illusion requires a large difference between expected and felt weight. Across all object pairs, perceived weight was well described by a model (R^2^ = .98) that includes a positive influence of both objects’ weights and the judged object’s density, but a negative influence of the other object’s density. Critically, the influence of both densities on perceived weight is strongly modulated by weight difference, being three times as large for zero / small weight differences than for large differences. Thus, it is only under the unusual conditions of typical SWI studies that density affects perceived weight to a substantial extent. Unlike existing models, that are inconsistent with our more comprehensive dataset, our descriptive model provides a quantitative, accurate and generalised account of weight perception for pairs of objects across various weight and size conditions.

**Author summary:** Knowing how heavy an object is allows us to grasp and lift it efficiently and without mishaps. Surprisingly, humans make systematic errors when judging the weight of objects of different size. For example, when two objects differ in size but have identical weight, the smaller object feels heavier. This is the ‘size-weight illusion’. The illusion is strong and reliable and occurs even when we know that the two objects actually have the same weight. The size-weight illusion demonstrates that the human perceptual system doesn’t work like a weighing scales, but instead takes into account other object features such as size or density, alongside weight. In this paper, we present a model that allows us to predict perceived weight in the size-weight illusion and across a wide range of objects of different weight / size / density, where previous models have failed.

## 1. Introduction

In the well-known size-weight illusion (SWI) (1), participants are typically presented with one large and one small object, apparently of the same material, but in fact of equal weight. On lifting, the smaller object is perceived as heavier than the larger one (Figure 1A). The illusion is robust, persisting even when the lifter knows that the objects have equal weight (2,3), and after grasping forces have adapted to the actual weight of the objects (4).

**Fig 1.**
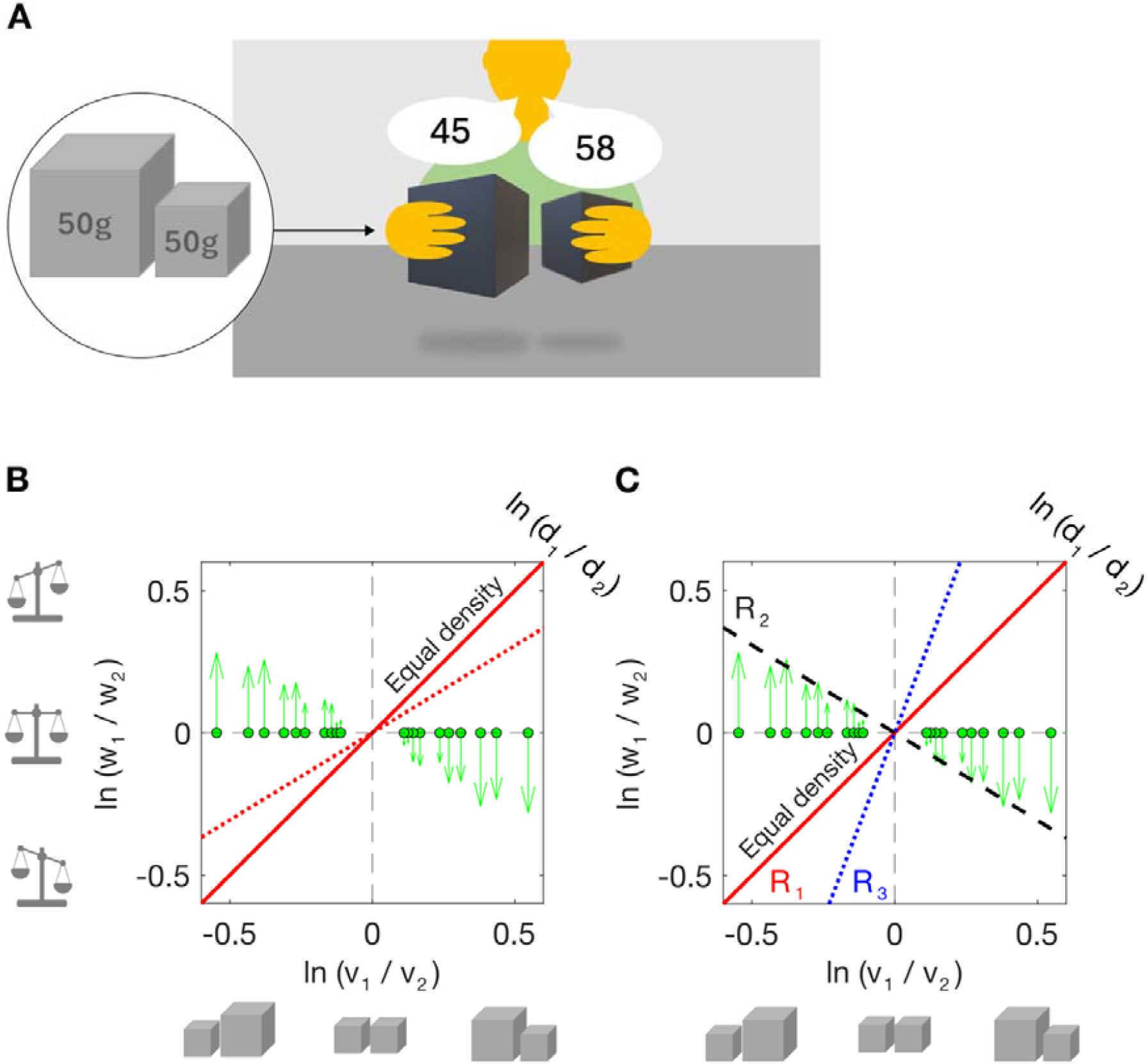
The size weight illusion. (A) Typical procedure for measuring the size-weight illusion (SWI). The subject lifts two objects of equal weight, but different size. The smaller object is perceived to be heavier than the larger one. (B) The SWI is ‘anti-Bayesian’. Stimulus pairs are plotted in a two-dimension stimulus space: log volume ratio (x-axis) and log weight ratio (y-axis). A value of zero indicates that the two objects have equal volume (or weight). The green markers show SWI stimuli (equal weight but different volume) and arrows show the empirical bias, i.e. the perceived weight ratio, for the same stimuli (data taken from the current experiment). The solid red line represents an equal density prior (log density ratio=0). The dotted red line indicates another plausible prior based on object statistics (5): the larger object is heavier but slightly less dense. In the SWI, perceived weight is biased away from these priors. (C) SWI and three hypothetical priors (R_1_ – R_3_) over log volume ratio and log weight ratio, as suggested by Peters and colleagues (6). Each prior corresponds to a different density / volume relationship. R_2_ (smaller is denser and heavier: dashed black line) would predict the SWI. R_1_ = equal-density prior (red line); R_3_ = smaller is less dense and lighter (dotted blue line).

Although the SWI has long been investigated as a means to understand human heaviness perception, its mechanism remains largely unexplained (for reviews, see (2,7,8)). Early descriptive accounts suggested that the SWI is caused by a contrast / repulsion effect, where the felt weights of the two objects conflict with those predicted by assuming equal density. Other accounts have suggested that the lifter confounds / combines felt weight with other physical features (generally density: (9–11); but also rotational inertia: (12,13)). Here we briefly describe existing models of weight perception that have been proposed to explain the SWI. We evaluate these models fully in section 5, ‘Modelling’, by determining how well they predict our data.

In ‘contrast’ or ‘violated expectation’ accounts it is noted that object size is used as a predictive feature for object weight (14–16). Smaller objects are expected to be lighter than larger objects, particularly when they are apparently made of the same material – that is, of equal density (see Figure 1B). Upon lifting, the smaller object is heavier than expected, and this disparity between expectation and sensory evidence is suggested to cause the illusion. Evidence for this conflict between expected and actual weight comes from the forces used to grip and lift the two objects. On initial lifts, participants use more force to lift the larger (expected to be heavier) object than to lift the smaller one (4,17). These misapplied forces have thus been suggested to drive the illusion: the larger object is lifted faster and more easily than expected, and vice versa for the smaller object, producing a mismatch between the expected and actual sensory feedback (17). However, it has been widely reported that grasping forces adjust to the weight of the object after a few lifts, while the illusion persists (2,4,18,19)).

Because the illusion is at odds with the perceiver’s prior expectations (that the smaller object should be lighter) the SWI has been labelled ‘anti-Bayesian’ (20,21). The Bayesian framework has successfully described numerous perceptual phenomena in terms of the integration (generally a weighted averaging) of prior expectations with current sensory cues (22–27). In the SWI, however, the Bayesian prediction is that the smaller object will be perceived as lighter than the larger object, given an assumption of equal density for the two objects. Figure 1 illustrates this: Figure 1B represents the stimulus space, with different possible combinations of two stimuli (their weight, volume and density log ratios). Two theoretically plausible, hypothetical priors are shown on these axes: (i) the larger object is heavier and same-density (solid line), or (ii) the larger object is heavier but slightly less dense (dashed line, reflecting measured object statistics (5)). Both of these priors predict the opposite of the SWI (green dots show stimuli, arrowheads give perceived log weight ratio, taken from current study). The SWI involves biases *away* from the theoretically sensible priors, hence the ‘anti-Bayesian’ label. The dashed black line in Figure 1C shows a hypothetical prior (smaller = heavier) that would predict the SWI. However, this prior is implausible, given environmental statistics in which size and weight are positively correlated.

Nonetheless, Bayesian models of the SWI have been proposed (6,28). Peters and colleagues (6) proposed a Bayesian model that includes three sub-priors over the relationship between the log volume ratio of two objects and their log density ratio (shown in Figure 1C). Each sub-prior represents a categorical density relationship; R_1_: objects have equal density, R_2_: smaller objects are denser and heavier than larger ones and R_3_: smaller objects are less dense (much lighter) than larger objects. Predicted biases in weight perception depend on the volume and density ratios of the stimuli and thus which sub-prior dominates perception. When a stimulus pair is encountered, the current sensory evidence (i.e., the sensed density and volume ratios of the two objects) is integrated with a prior over the density relationship of the two objects, which incorporates the three subpriors R_1-3_. The sub-prior that dominates perception, i.e., that dictates the direction of any perceptual bias, depends on the current sensory evidence. When a typical SWI stimulus pair (e.g. log weight ratio = 0, log volume ratio = 0.5) is encountered, the ‘smaller is denser and heavier’ prior (R_2_ in Figure 1C) dominates, resulting in a size-weight illusion. It is not clear whether the model can account for weight perception more broadly as it was only evaluated on pairs of equal-weight objects (i.e., typical SWI stimuli).

Wei and Stocker (29,30) have suggested that contrast effects analogous to the SWI, i.e. biases away from a prior, can be predicted by Bayesian models incorporating efficient coding of stimulus properties (31,32). They note that when efficient coding maximises Fisher information (i.e., to reflect the stimulus distribution in the environment), this results in likelihoods that are systematically skewed with a long tail away from the peak of the prior. If a symmetric loss function is also employed (selecting the mean rather than the maximum of the posterior: L2 loss), estimates will be systematically biased away from the prior, relative to the peak of the likelihood (‘likelihood repulsion’), see ‘Modelling’ section for more detail. Recently, Bays (28) has proposed Bayesian models of the size-weight and material-weight (33) illusions that incorporate efficient coding and priors reflecting natural statistics. In the case of the SWI, the prior reflects a positive association between size (volume) and weight. However, whilst Bays’ model provides predictions of perceived weight across the whole volume-weight space, it has only been tested on Peters and colleagues’ (6) dataset.

Other authors suggest that we are unable to fully separate density and weight (34), or proactively combine them (11). Under these models, denser objects are systematically perceived as heavier. Indeed, weight and density estimation appear to be closely related: the difference in estimated weight increases with increasing density difference (34); objects of greater weight are reported as denser (35), and weight discrimination is improved when density varies in the same direction as weight (36). Within typical SWI stimuli, weight is kept constant such that density is negatively correlated with volume. Thus, size contributes to the illusion because it is used to infer density (9,11,37).

Wolf and colleagues (10,11) proposed a cue-integration model in which perceived weight is a weighted average of two heaviness estimates, one from the object’s felt weight and one derived from the object’s inferred density. Because denser objects tend to be heavier, the authors suggest that it is adaptive / Bayesian to combine estimates of weight and density when estimating weight, to obtain more reliable (less noisy) weight estimates. It is not clear, however, why we should combine weight and density information in this way, when volume information is available to condition the density-weight relationship. Not only is density a poor cue to weight (when not conditioned on size) but the two estimates will also have correlated noise (thus negating some benefits of integration).

In general, the focus on density rather than volume in existing explanations of the SWI seems somewhat counterintuitive. After all, weight and volume are directly sensed, whereas density must be inferred from the other two. Nonetheless, previous models and the long-standing ‘violated expectations’ narrative have focussed on density. The expectation that adjacent objects, apparently comprised of the same material, will have similar densities seems reasonable.

It is well known that the SWI magnitude increases with volume (density) difference (6,8,34,38,39) and this relationship is predicted by all of the above models. SWI demonstrations tend to use fairly large, obvious differences in object volume, in order to obtain a reliable illusion. However, under small conflicts between expected weight (based on size, under an assumption of equal density) and direct sensory evidence, these estimates might be integrated. We know that integration occurs for small conflicts but breaks down under conditions of large conflict (40,41), but this has not been tested in the context of the SWI. If integration (the opposite of the SWI) occurs under small conflicts, while the SWI only occurs with large conflicts, this would be broadly in line with the idea that the SWI is driven by violated weight expectations. Yet, it should be noted that an absence of integration does not directly account for the contrast effect observed in the SWI. Integration under small conflicts would be consistent with Peters and colleagues’ model (6), with integration driven by their equal density sub-prior (see above). Under small density differences (log density ratios close to 0) the sensory evidence is close to the equal density sub-prior, and the model thus predicts integration, i.e. attraction towards equal density.

However, integration under small density differences would be at odds with models in which weight and density are confounded / integrated in general (10,11). Furthermore, it is unclear whether a conflation of weight and density can account for perceived weight in all situations (such as when a smaller object is less dense and therefore much lighter than a larger one), or only in the specific and non-accidental conditions of SWI stimuli, when weight is held constant and volume varies.

Here we asked: can we develop a generalised model of weight perception for varied combinations of size, weight and density? To this end we measured perceived weight for pairs of cubes from a large stimulus set in which volume, and / or weight, and / or density were varied in small steps. Stimulus pairs could have equal or different density, and if different, the denser cube could be larger or smaller than the less dense cube. Size and density were uncorrelated in the full set to prevent learning of unusual size-weight associations. For each stimulus pair, participants viewed and lifted both cubes before reporting each of their weights (i.e., magnitude estimation – see Figure 1A). To preview the main findings: 1) The SWI occurred even at very small size (density) differences, and thus does not rely on large conflicts. 2) Perceived weight was positively correlated with the true weight of both the judged and other stimulus, in addition to the judged object’s density. 3) The effect of both objects’ densities on perceived weight was strongly modulated by the weight difference between the two objects: three times larger at small / zero weight differences than at large weight differences. In fact, the other object’s density had an effect only when the two objects had similar weight, when it was negatively associated with the judged object’s perceived weight. Thus, density affects weight perception, but this effect is much larger in the specific and peculiar conditions of typical studies of the SWI. We provide a model (R^2^ = .98) that captures biases in perceived weight in the SWI and across the full stimulus set. In addition, we compare this model to existing models of the SWI, and to a novel efficient-coding Bayesian model of the SWI.

## 2. Materials and Methods

### 2.1 Participants

Thirty naïve participants (age *M* 21.4, *SD* 5.6 years) from the University of Southampton Psychology student community took part in the experiment; undergraduate students were compensated for their participation with course credits. All reported normal or corrected-to-normal eyesight and no upper limb impairment. The experiment was approved by the University of Southampton Psychology Ethics Committee (No 47316.A1–A3); all participants gave written prior informed consent.

### 2.2 Stimuli

Stimuli were 15 plastic cubes (3D-printed in black polylactic acid (Figure 2A–B). Each cube was filled with sand, polystyrene pellets, and / or brass to create the desired volume, weight and density values for the different stimulus sets, depicted in Figure 2A. The centre of mass of the cubes remained at the geometric centre and the filling did not move inside the cubes during lifting. A panel concealed at the bottom kept the appearance homogeneous. Weight was measured with a 0.1 g resolution.

**Fig 2.**
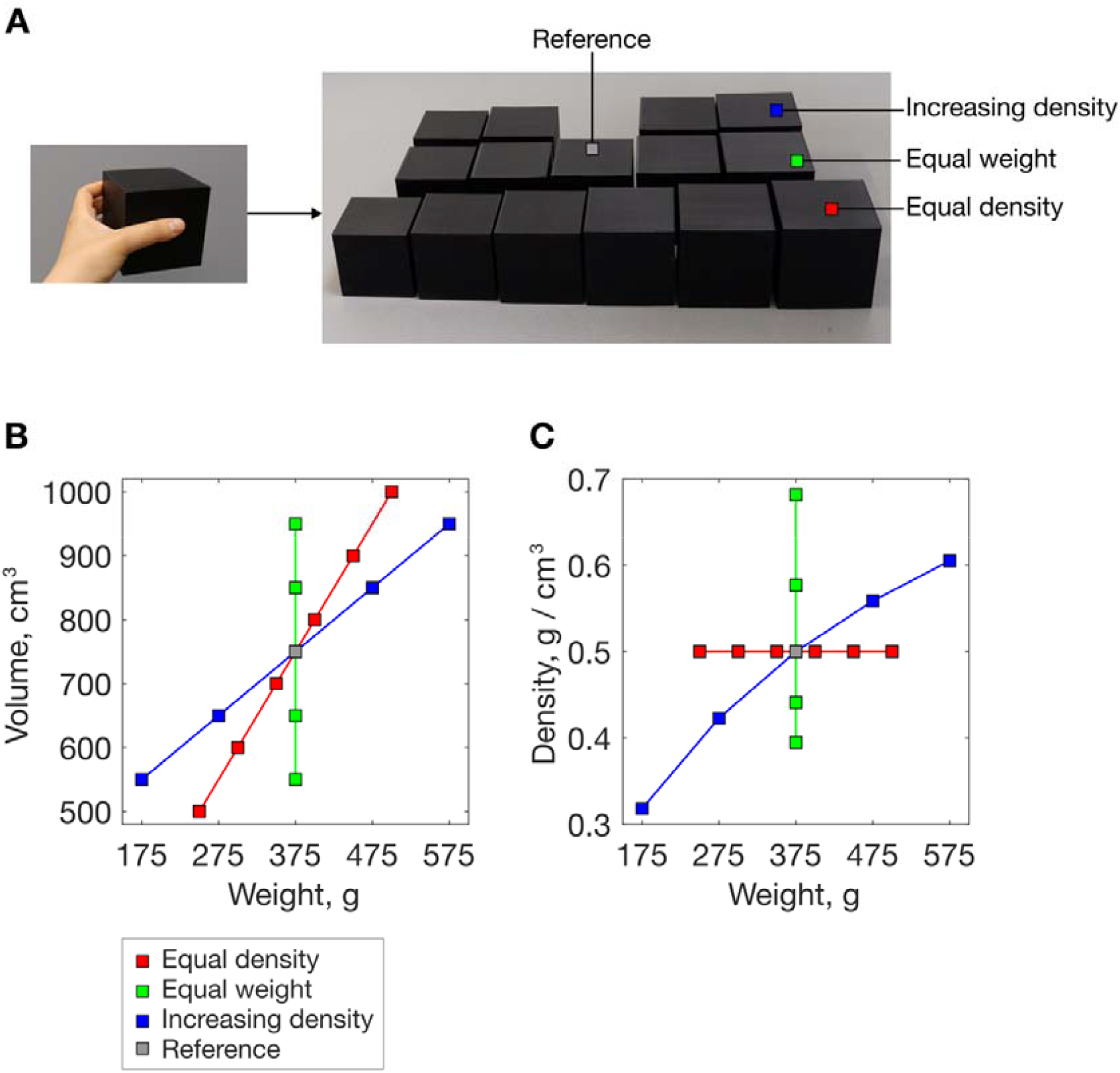
Stimulus set. (A) Full stimulus set, with 500 cm^3^ stimulus for scale on the left. (B) Stimulus volume as a function of weight. (C) Stimulus density as a function of weight.

The complete stimulus set included three subsets (Figure 2B–C), corresponding to different volume / weight relationships: 1) Equal density (red): weight increased linearly with volume (as for ‘normal’ same-material objects); 2) Equal weight (green): density decreased with increasing size (typical SWI stimuli); 3) Increasing density with increasing size (blue): both weight and density increased with size. The middle cube (volume, weight, density: 750 cm^3^, 375 g, 0.5 g / cm^3^) was common to all three subsets. Volumes within each set increased in 100 cm^3^ steps, with the addition of 50 cm^3^ steps below and above the middle stimulus in the fixed density set. Minimum (maximum) volume, weight, density differences across any stimulus pair were: 50 (250) cm^3^, 25 (200) g, 0.02 (0.18) g / cm^3^, respectively. Density ratios for stimulus pairs were in the range 0.47–0.97 (less dense / denser stimulus), 0.58–0.89 g / cm^3^ for equal weight (SWI) stimulus pairs. This contrasts with previous work, where smaller ratios (bigger differences) in density have been explored (largest ratios for classic SWI pairs: 0.67 (42), 0.64 (43), 0.51 (44); a larger ratio of 0.87 was tested by Wolf and colleagues (11) using same-volume pairs).

### 2.3 Task and trials

Stimuli were presented in pairs. In each trial, participants simultaneously grasped and lifted both cubes in full visuo-haptic conditions, and verbally reported both objects’ weights using magnitude estimation (as in Figure 1A). The paired presentation (common in SWI experiments), alongside magnitude estimation, allowed us to model the perceived weight of each object in addition to the pair’s difference in perceived weight or ratio of weight estimates. Thus, we could evaluate models that consider judgements of absolute perceived weight (11,28) in addition to models that only consider the relative perception of stimulus pairs (6).

To reduce variability and response drift that can be associated with magnitude estimation, we provided a reference object at the beginning and at regular intervals throughout the experiment. The reference object was part of the stimulus set (see below, 2.5).

Experimental trials included all possible stimulus pairs (same- and different-subset pairs), excluding pairs of identical stimuli. This gave 105 unique pairs and 133 pairs in total (trials including the middle / common stimulus were repeated due to a coding error). Each stimulus pair was presented twice, with left and right position switched between the two trials (266 trials in total). Trials were randomised for each participant.

### 2.4 Setup

The participant and the experimenter sat at opposite sides of a table covered in thick felt to muffle auditory cues to the objects’ weights. Two numbers (1, 2) marked the positions of the stimuli on the table (centre-to-centre distance between stimuli, centre of stimuli to table edge: both 20 cm) and served to remind participants the order in which to report their estimates. A frame-mounted black curtain was used to control stimulus presentation, such that the participant only ever saw the two stimuli of the current trial. A desktop computer running MATLAB (45) was used to randomise stimulus presentation and record participant responses.

### 2.5 Procedure

The experiment was completed in two identical laboratory sessions on separate days, each lasting 60–90 minutes. At the beginning of each session, participants were told that their task would have been to lift the two cubes and provide an estimate of their weight, in arbitrary units. They were presented with the middle cube of the stimulus set (750 cm^3^, 375 g, see Figure 1A-C) as the reference stimulus, and told that this had a weight of ‘50’ (arbitrary units), and that they will have to judge the weight of each cube in reference to this (‘If you think that one cube weighs half of the reference weight you will say 25. If you think that it weighs double the reference weight you will say 100. You can use any number, just use the number that best represents the weight of each cube’). The reference was presented again every 20 trials (in addition to featuring in some of the stimulus pairs). On each trial, participants were instructed to look at the stimuli while lifting. Exploration mode was not restricted: participants were free to grasp and lift the cubes as they thought best; they were not restricted to use index and thumb only. This was done to provide the most reliable size estimates (11). The number of lifts within each trial was also not restricted. Trials were not timed, but participants were encouraged to give an answer if they waited more than 30 seconds (approx.).

## 3. Results

Participants’ magnitude estimates were transformed into grams and then normalised to remove individual differences in how the response scale was used: Each participant’s estimates were multiplied by the ratio of their individual grand mean across stimuli to the grand mean across all participants (thus, after normalisation, each observer had the same mean response). Outliers per condition were then removed iteratively: for each object within each stimulus pair, values more than 5 SDs, then 4 SDs, and 3.5 SDs from the mean across observers were removed (65 datapoints in total, or 0.4%). Data were re-normalised after each iteration. All further analyses were conducted on the resulting dataset.

Weight estimates for within-subset pairs (i.e., both objects from the same subset) are shown in Figure 3, alongside model fits (see below, Equations 1–2). Data have been collapsed across repetitions within participants and *SE* bars show inter-observer variation. We present the perceived weight of the judged object (i.e., the object within each pair whose weight is being reported) in terms of its own properties (weight / density) and the properties (weight / density) of the other object in the pair. Each object in turn was considered the judged object.

**Fig 3.**
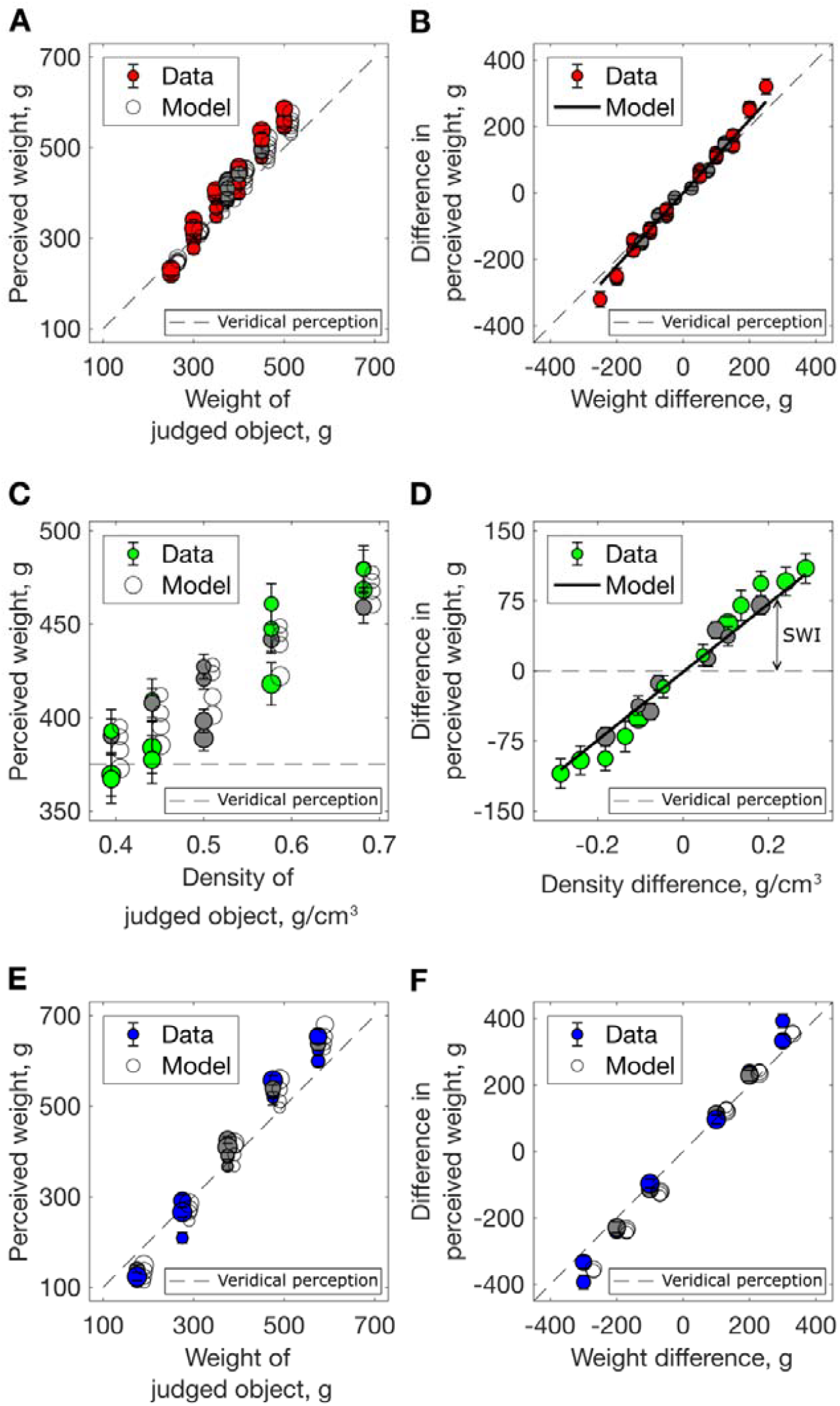
Results for within-subset stimulus pairs. Perceived weight (A, C, E) difference in perceived weight (B, D, F) for within-subset stimulus pairs, averaged across participants, error bars show ± 1 SEM. Solid lines / open symbols show the model predictions (described below, Equations 1 and 2). The dashed lines indicate veridical perception. Grey markers indicate pairs that included the reference stimulus (either as judged or other object). (A–B) Equal density pairs: (A) Weight estimates as a function of stimulus weight; symbol size indicates the weight of the other stimulus in the pair. (B) Data re-expressed as difference in perceived weight, as a function of weight difference. (C, D) Equal weight (SWI) pairs: (C) Weight estimates as a function of stimulus density; symbol size indicates the density of the other stimulus in the pair. Figure A (panel A) in S1 Supporting information shows weight estimates as a function of stimulus volume. (D) Data re-expressed as difference in perceived weight as a function of density difference; symbol size indicates the average object density. Figure A (panel B) in S1 Supporting information shows the difference in estimated weight as a function of volume difference. (E, F): Increasing density pairs: (E) Symbol size indicates the other stimulus’ weight; (F) Data re-expressed as difference in perceived weight, as a function of weight difference; symbol size indicates the average object density.

For equal density, i.e. ‘normal’ object pairs (Figure 3A–B), weight estimates were highly correlated with the true weight of the judged object, as expected. However, they were also positively correlated with the weight of the other object (denoted by symbol size in Figure 3A; see the vertical spread corresponding to weight of other object); note that this is an averaging, rather than a contrast effect. The difference in perceived weight between the two objects was linearly related to stimulus weight difference (Figure 3B; data are symmetrical in all difference plots (B, D, F) as each object in each stimulus pair serves as ‘judged’). We asked participants to estimate the individual weights of the two objects in each pair (not to estimate the difference between their weights). However, a control experiment confirmed that these two tasks produce mutually consistent data (see S1 Supporting Information).

Weight estimates for the equal weight (SWI) pairs are shown as a function of the judged object’s stimulus density (x-axis) and the other object’s density (symbol size, Figure 3C). In line with the SWI, perceived weight was positively correlated with object density: denser (and smaller) stimuli were perceived as heavier (overall positive slope of the data). Perceived weight was also negatively correlated with the other object’s density, the judged object appearing heavier when judged next to less dense (larger) objects (see the vertical spread of data points in 3C). The data are re-expressed as difference in perceived weight as a function of density difference in Figure 3D. The magnitude of the SWI (i.e. deviation from veridical perception) increases with density difference, as expected, but still occurs with small density (or volume) differences. Thus, we find no evidence for integration of expected and sensory weight estimates under small conflicts.

Lastly, in the increasing density pairs (Figure 3E), weight estimates were positively correlated with the object’s weight, but also correlated with the other object’s weight (as in the equal-density pairs; see vertical spread of data in 3E). Figure 3F shows that the difference in perceived weight increases linearly with true weight difference. In this set, weight (or weight difference) is correlated with density (or density difference); the effect of density contributes to the increased slope of the data, relative to veridical perception (identity line). All of the effects described above were observed regardless of whether or not the reference stimulus was included in the stimulus pair (grey markers in Figure 3).

Figure 4A summarises results for all trials, including across-subset pairs (i.e. objects within a pair are from different stimulus subsets), in a format that illustrates the variable influence of density on perceived weight. For every stimulus pair (represented by a dot), the difference in perceived weight (y-axis) is presented as a function of stimulus weight difference (x-axis) and stimulus density difference (colour). Lines show the model fit (see Equation 2 below). The overall slope of the data shows the strong relationship between true weight difference and the difference in perceived weight. The vertical spread of the different colours shows the effect of density difference on the difference in perceived weight. The two are not independent: there is a central ‘bulge’, where the effect of density is larger (more vertical spread). In other words, the effect of density (or density difference) on perceived weight (or difference in perceived weight) is strongly modulated by stimulus weight difference, being much greater for small / zero weight differences. Figure 4B summarises the influence of the two objects’ densities on perceived weight, as captured by the model’s density coefficients (see below). The conflation of weight and density is dramatically increased under the conditions of typical size-weight stimuli, where weight difference is zero.

**Fig 4.**
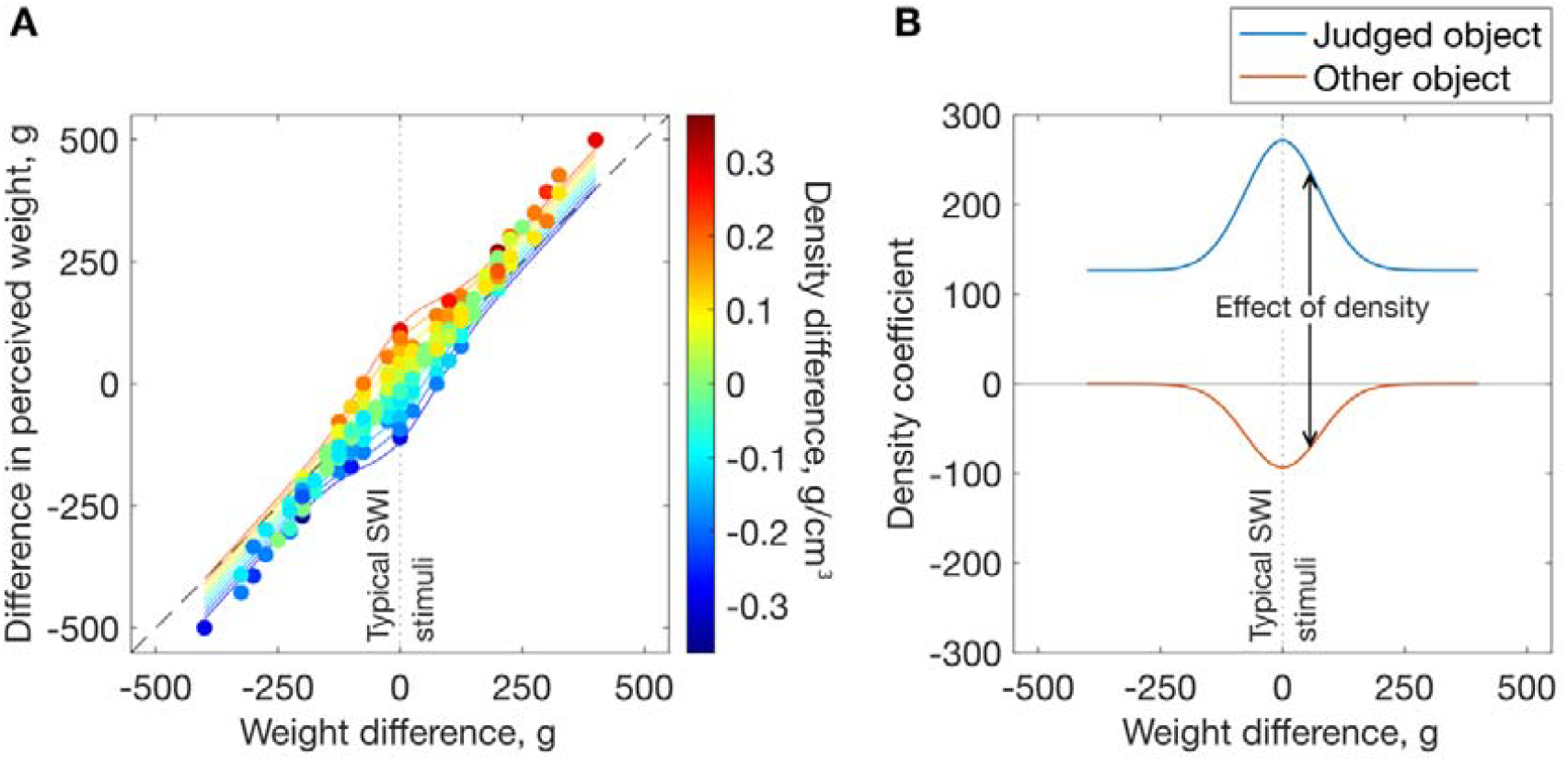
Effect of density. (A) Difference in perceived weight as a function of weight difference and density difference in the full dataset. Markers show the data, solid lines show the model predictions (see Equation 2) at equally spaced density differences. The vertical spread of the data with density difference (marker colours) shows the effect of density in typical SWI stimuli (i.e., equal weight: vertical dotted line) and across the full set. The dashed diagonal line shows veridical perception. (B) The influence of density (i.e., density coefficients) as a function of weight difference in the model.

## 4. Comparison with existing literature

The basic SWI (smaller / denser objects are perceived as heavier than their larger, equal weight counterparts) has been confirmed via a wide array of experimental paradigms. Stimuli have been presented singly (e.g., (46,47)) or in pairs. Paired stimuli are lifted sequentially using the preferred hand (e.g., (4,6,48)) or simultaneously, one per hand (49–52); simultaneous and sequential lifting yield SWIs of similar magnitude (53). Studies that measure fingertip forces (e.g., (4,48,54)) usually employ sequential lifting; only one force-sensitive handle is needed, and any hand-dependent force differences are eliminated. A few studies involving heavier objects used bimanual (38,55) or even team lifting of single objects (56,57). Plaisier and Smeets (58) elicited the SWI without any lifting: participants applied lateral force to objects suspended on strings.

The vast majority of SWI studies have employed magnitude estimation; subjects ascribe a value to each object’s perceived weight. In some, no explicit reference or scale is provided, as in absolute magnitude estimation (e.g., (47)) or when using bounded estimates (e.g., 1–10; (54)) with the requirement to adhere to a single scale throughout sometimes made explicit (e.g., (11)). Without an explicit reference scale, observers must rely on an established scale (e.g., grams, learnt from multiple reference objects), or use an idiosyncratic scale defined by the first stimulus and their response to it. These strategies may be prone to noise (uncertainty) and drift. For bounded estimates, subjects may be forced to change their scale if an ‘out of range’ stimulus is encountered. Other studies, therefore, provide an explicit reference object. This object may be part of the experimental stimulus set (e.g., the current study; (39,59) or a separate object (46). Peters and colleagues (6) used a moving reference scale: one object within each pair was labelled as ‘10’ on each trial. Only a few studies have rejected magnitude estimation in favour of a two-alternative, ‘which object is heavier (lighter)’ task (e.g., (20,50,51).

Despite the variety in methodology, some commonalities emerge within and across studies: (i) a SWI is observed, (ii) larger density differences result in a larger SWI, and (iii) the availability of haptic size information results in a larger illusion (for a meta-analysis, see (8)). Within the data from our equal-weight trials (Figure 3C-D) a clear SWI is present, and its magnitude increased with increasing density differences. Importantly, however, the present study goes beyond these demonstrations of the classic SWI: we place it in the broader context of weight perception and systematically explore how the weights and densities of objects interact to predict perceived weight (as highlighted by Figure 4). The following section evaluates existing and new models of weight perception, with respect to how well they account for this richer dataset.

## 5. Modelling

### 5.1 Model of perceived weight

Our model captures perceived weight 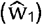 for all stimulus combinations, as a function of the weight and density of the judged (w_1_, d_1_) and other object (w_2_, d_2_). We note that in any task involving magnitude estimates we cannot, as experimenters, know how a participant’s internal perceptions map on to their reported judgements. This is the classic El Greco problem of perception: perceptual biases can affect perception of all objects, including reference objects (60,61). For this reason, we do not focus on absolute perceptual biases (i.e., stimulus X is perceived as Y units heavier than its true weight). Instead, we focus on how estimated weight is *modulated* by the properties of the stimulus and the context (i.e. the other object within the pair), as captured by the model.

Our model is presented as a schematic in Figure 5A, which shows the 5 components of the model, and described formally by Equation 1; perceived weight is positively correlated with both objects’ weights, and the judged object’s density. When the two objects’ weights are similar (as in the SWI), the two objects’ densities have an additional influence (dictated by a Gaussian over weight difference). The model (Equation 1) explains 98% of the variance in the full dataset:

**Fig 5.**
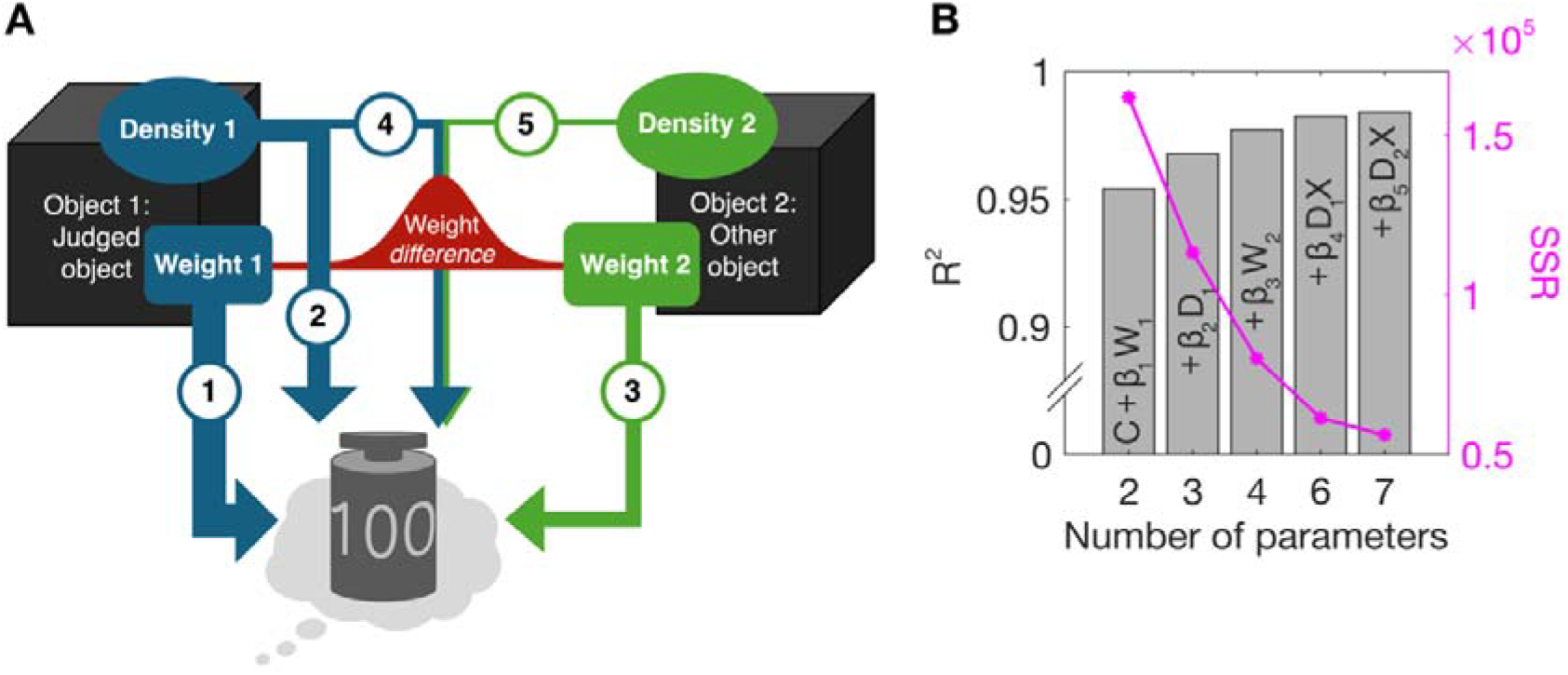
Model of perceived weight. (A) Schematic of the model’s 5 components. The perceived weight of an object is positively correlated with both objects’ weights (components 1 and 3), and the judged object’s density (component 2). When the objects’ weights are similar (captured by a Gaussian over their difference) the two objects’ densities have an additional influence (components 4 and 5). The importance of each component is conveyed by the thickness of the corresponding arrow. (B) Variance in perceived weight explained by the addition of each component to the constant in the model (bars) and corresponding SSR (magenta lines and markers) (C). The order of components follows the maximum increase in R^2^ with each addition (see Table A in S2 Supporting Information). X =.

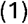

Figure 5B shows how the model fit improves with each additional component (i.e., components 1-5, as in Figure 5A) of the model.

Our model was fitted to perceived weight estimates for single objects (_1_). Accordingly, the difference in perceived weight (_1_ – _2_) between two stimuli can be predicted via the following equation:

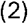

Perceived weight is positively correlated with the object’s weight (ß_1_ = 1.23) and its density (ß_2_ = 126.43), and with the other object’s weight (ß_3_ = 0.13). However, when the two objects have similar weight, the effect of the object’s density is magnified, and the other object’s density also has an effect. This interaction between weight difference and the effect of density is captured by a Gaussian centred on a weight difference of 0. The density of the judged object positively influences its perceived weight (ß_4_ = 27814), whereas the other object’s density has a negative influence (ß_5_ = −17919). Sigma for the Gaussian component is 76.44. Figure 4B shows the influence of both objects’ densities as a function of weight difference: the combined influence of the two densities on perceived weight is approximately three times larger for zero weight differences than for large weight differences. All analyses were done in MATLAB (45) and optimal coefficients (minimising the summed squared residuals (SSR) in the pooled data) were identified via gradient descent. To prevent over-fitting, we used leave-one-out cross-validation on the normalised dataset, leaving out each participant in turn. This method ensures that every component in the model improves its ability to predict data from new observers (62–64). Alternative candidate models are presented in Table A in S2 Supporting information alongside model comparison statistics. Given that volume and weight are directly sensed, whereas density must be inferred, we also tested models including the objects’ volumes (v_1_, v_2_) instead of density as predictors, but these were inferior, see Table B in S2 Supporting information.

Whilst our model provides an excellent quantitative account of perceived weight, it does not explain *why* our sensory system computes perceived weight in this way. Below we explore whether a Bayesian model incorporating efficient coding might provide a more theoretically appealing account of biases in weight perception. We then compare our models to those previously proposed to explain biases in weight perception.

### 5.2 Efficient coding Bayesian ratio model

Traditionally, Bayesian models have been employed to account for perceptual biases towards a prior (i.e. towards the most frequent or probable stimuli). Examples in the literature include biases for light-from above (65), isotropy assumptions when interpreting foreshortening (40), global convexity of surfaces (66), and ‘slow and smooth’ motion (67). However, as described by Wei and Stocker (29,30), Bayesian models incorporating efficient coding of stimulus features can also predict repulsion biases, i.e. biases *away* from the prior. Given that the SWI (and not integration) is observed even for very small density differences, we asked whether such a model might provide a quantitative, Bayesian model of the SWI as a ‘repulsion’ effect, where perception is biased away from a prior for equal density. Wei and Stocker (29,30) show that the expected perceptual bias for a stimulus value from a stimulus distribution can be predicted from the prior, p() (in fact from the slope of its squared reciprocal), as in Equation 3:

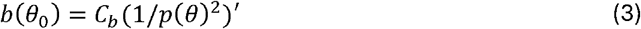

where *C*_*b*_ is a constant.

We present a simple model that includes a prior over the relationship between two objects’ weights and volumes: i.e. the two dimensions of log weight ratio and log volume ratio, as depicted in Figure 1B. Such a prior, if aligned with the log weight ratio = log volume ratio unity line (solid red line in Figure 1B), represents the assumption that the two objects have equal density. Perceptual biases *away* from this prior would be consistent with the ‘contrast’ or ‘repulsion’ explanation of the SWI that has prevailed for decades (2,7). Thus, similar to Bays (28), we use the work of Wei and Stocker (29,30) to model the seemingly anti-Bayesian bias in the SWI as repulsion away from the prior. Unlike Bays’ model, our ratio model predicts biases in perceived weight ratio for pairs of objects, given a prior over their weight and volume ratios. Bays’ model fits the perceived weight of single objects (see below, 4.3). For pairs of objects, considering ratios of weight, volume and / or density (rather than their differences) is a natural choice: any two of these ratios define the third, as (w_1_/w_2_) / (v_1_/v_2_) = d_1_/d_2_. Taking the log ratio gives a useful symmetry: log(w_1_/w_2_) = −log(w_2_/w_1_); allowing us to predict the same magnitude of bias, irrespective of which object in a pair is designated as object 1. To fit the model, we calculated the ratio of weight estimates on each trial for each participant. Like Peters and colleagues (6), we did not ask participants to provide ratio estimates, as we expected they might be unfamiliar with using ratios. Our prior is defined by a mixture of three bi-variate Gaussians centred on [0, 0], i.e. equal weight and volume (Figure 6A). This mixture-model allowed us to explore variations in the shape of the prior; there is no reason to assume that it should be a perfect bivariate Gaussian. Model parameters and model comparisons given priors of different complexity are provided in Table C in S2 Supporting Information.

**Fig 6.**
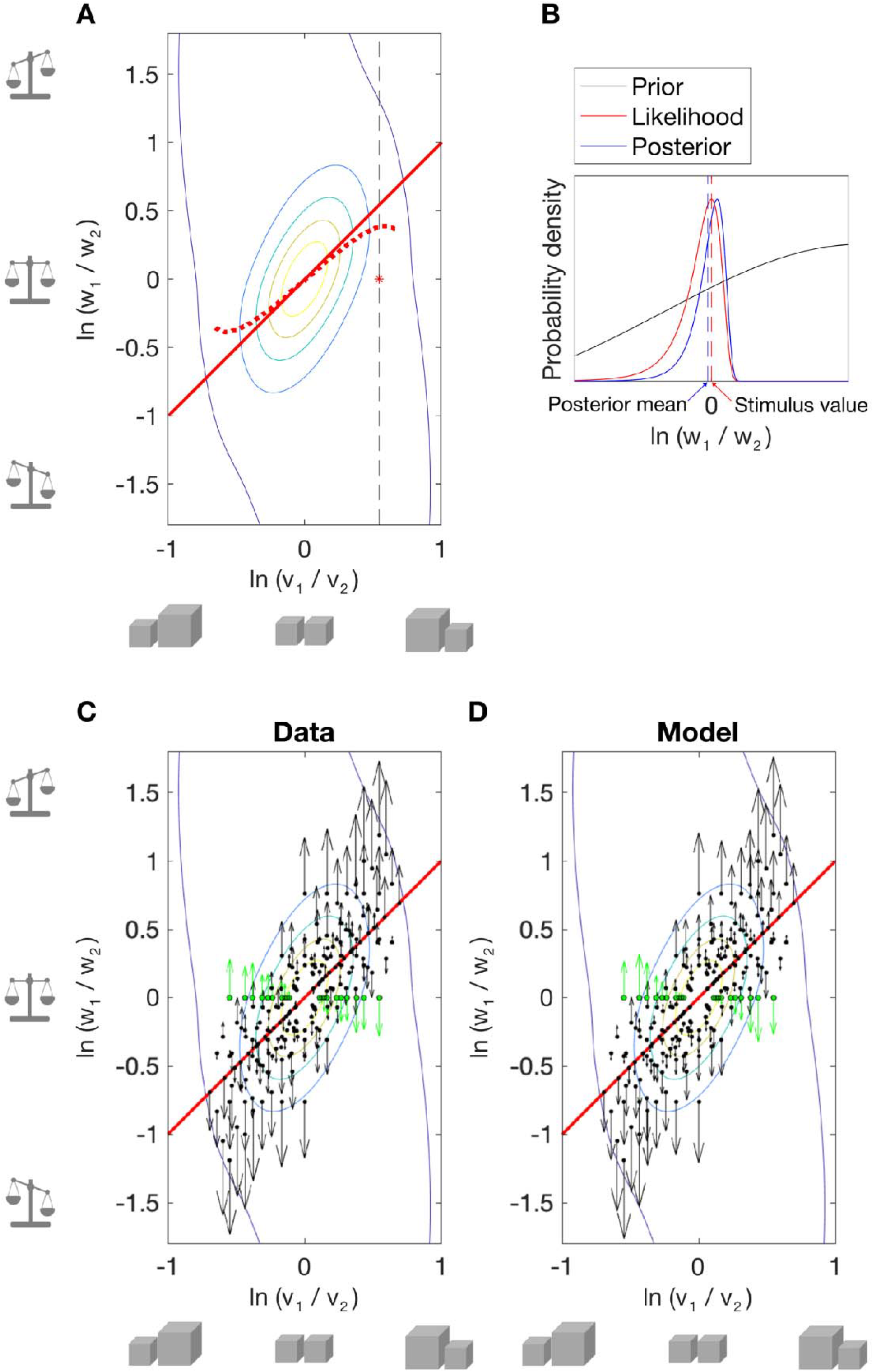
Efficient coding Bayesian ratio model. (A) Bi-variate prior over log weight and log volume ratio, optimised to minimise errors in predicted log weight ratio. Red asterisk and dashed black line indicate an example stimulus condition and corresponding slice through the prior that are shown in part B. The solid red line is the identity line (i.e. equal density), and the dashed red line shows the orientation of the fitted prior (plotted over the stimulus range) as defined by its peak values. (B) Likelihood repulsion predicted by Wei and Stocker (29) for one stimulus condition with equal weight, unequal volume stimuli. Schematic of a median (unbiased / noise-free) observer likelihood (red) and posterior (blue) for an example stimulus pair (red asterisk in A), given the 1D slice through prior (solid black line). Efficient coding of stimulus properties results in skewed likelihoods with a long tail away from the peak of the prior. The posterior is also skewed. When a symmetric loss function is chosen, e.g. the mean of the posterior (L2, blue dashed line), a bias away from the prior is predicted, i.e. the posterior estimate is shifted away from the prior peak, relative to the peak of the likelihood. Measurement noise (expected to be skewed in stimulus space) further increases the predicted repulsive bias (29). (C-D) Biases in perceived log weight ratio. Contour lines show the prior; black dots show the stimulus pairs; arrows show the magnitude and direction of the bias for each pair, with SWI conditions in green; the solid red line shows equal density. (C) Empirical biases; (D) predicted biases.

Similarly to Bays (28), we employ the simplifying assumption that perceptual estimates of volume ratio are relatively precise, and thus not significantly affected by the prior. Indeed, there is no clear evidence that objects with unexpected weights (e.g. typical SWI objects) cause biases in *volume* perception (i.e., a ‘weight-size illusion’ (68,69)). Thus, we estimated the expected bias in perceived log weight ratio by taking a slice through the prior at the true value of log volume ratio (e.g., vertical dashed line in Figure 6A) and finding the slope of this 1D prior with respect to log weight ratio (Figure 6B). If volume ratio judgements were also available, one could drop the simplifying assumption, and simultaneously fit biases in log weight ratio and log volume ratio. The resultant fitted prior would have a slightly different shape.

We fitted the model via gradient descent to optimally predict biases in perceived log weight ratio, averaged across participants, for all unique stimulus pairs (N = 105). Figure 6A shows the best fitting prior, whose orientation is close to an equal density prior (solid red, identity line), but is in fact rotated slightly towards larger objects being heavier and slightly less dense (dashed red line) in keeping with the object statistics reported by Peters and colleagues (5). The empirical biases in perceived log weight ratio are shown in (C) and those predicted by the fitted prior are shown in (D). The model accounts for perceived log weight ratio well, with 12 parameters (R^2^ = .988; see Table C in S2 Supporting Information). However, goodness-of-fit is poorer than our descriptive model (Equation 1, 7 parameters), when parameters are optimised to minimise the SSR in perceived log weight ratio (R^2^ = .991, see also cross-validation errors, Table A in S2 Supporting Information). Nonetheless, this efficient-coding Bayesian model provides a theoretically appealing account of the SWI; biases in perceived log weight ratio are minimal close to the d_1_=d_2_ identity line (solid red lines in Figure 6 C-D) and increase with log density ratio, broadly in line with repulsion away from an equal-density prior. Indeed, biases in log weight ratio correlate quite strongly with log density ratio (R^2^ = .691).

Tempting as it is to break out the champagne and toast our Bayesian brains, this may be premature. Closer inspection yields some difficulties with this model. First, the model is agnostic regarding how the bias in log weight ratio is split across the two objects. A reasonable assumption is that the prior has equal and opposite effects on the perceived weights of the two objects. However, this is not observed (see Figure B in S2 Supporting Information); there is only a weak correlation (R^2^ = .283) between biases in estimates of log weight ratio and biases in estimates of log w_1_ and log w_2_. In fact, biases in estimates of log w_1_ and log w_2_ are largely independent of the two objects’ relative densities (R^2^ = .215), in contrast to what we would expect given repulsion from an equal density prior. The pattern of bias in log weight ratio is fairly well approximated by a model that predicts each object’s perceived weight from its weight and density, that is, independently of any interaction between the two objects (R^2^, biases in log weight ratio =. 847 see Table A in S2 Supporting Information). This suggests that, although empirical biases are consistent with repulsion from an equal-density prior (Figure 6B), this might not be the underlying mechanism. It is worth noting that any pattern of bias can be explained with a complex enough prior. Observed and predicted biases in judgements of log w_1_ and log w_2_ are shown in supplementary Figure B in S2 Supporting information.

Second, a key feature of Wei & Stocker (29,30) is that the model gives predictions for both biases *and* reliability (discrimination thresholds or estimate variability). According to the efficient coding model, more frequently observed stimuli are represented more precisely, such that discrimination thresholds are proportional to the reciprocal of the prior (see Efficient-coding Bayesian ratio model in S2 Supporting information, for fitting of response variance). At the individual observer level, our data provide poor estimates of discrimination (only two estimates per pair). However, at the group level, a systematic pattern is clear (see supplementary Figure C in S2 Supporting information). Empirical response variance is well predicted by the absolute log weight ratio (R^2^ = .532); observers’ judgements of relative weight are most reliable when the two objects’ weights are similar. Log volume ratio adds little explanatory power (R^2^ = .557, for regression model predicting response variance from log weight and log volume ratios). Predictions from the efficient coding Bayesian model follow the opposite pattern: modelled response variance depends primarily on log volume ratio (R^2^ = .765) and far less on log weight ratio (R^2^ = .86, for regression model including both log volume and log weight ratios). In fact, the empirical pattern of response variance is better predicted by log weight ratio than by the modelled pattern of response variance (R^2^ = .532 vs R^2^=.444). For a variety of perceptual tasks, Wei & Stocker’s approach explains both the pattern of bias *and* the pattern of response variance (or discrimination thresholds) with a single fitted prior (30). Unfortunately, that is not the case for perceived weight, and is therefore unlikely to represent the underlying mechanism of the SWI. (We also considered, and rejected, Bayesian models of perceptual bias involving a 2D prior over *differences* in weight and *differences* in volume or density, see ‘Efficient-coding Bayesian ratio model’ in S2 Supporting information).

### 5.3 Other models

Here we explore how well existing models of weight perception by Bays (28), Peters and colleagues (6), and Wolf and colleagues (11) (see Introduction) explain our data.

Bays (28) proposed a Bayesian model of the SWI, which, as above, predicts biases away from the prior, given efficient coding and an L2 loss function (29). However, rather than considering object pairs, Bays’ model employs a bivariate Gaussian prior over the perceived weight and volume of single objects, i.e., a positive association between weight and size.

Supplementary Figure D in S2 Supporting information shows the model’s fit to our data. We extended the model to incorporate a mixture of Gaussians prior (Table D in S2 Supporting information for model parameters and model comparisons). Whilst the model captures the mean bias for each stimulus reasonably well, it cannot capture the way in which perceived weight is modulated by the other object in the pair, nor is it consistent with the pattern of response variance (see Figure E in S2 Supporting information). As noted above, Bays’ (28) model predicts biases in the estimated weights of single objects, given a prior over the properties of single objects. However, one might suggest fitting Bays’ model directly to the log ratio of the perceived weights of object pairs (i.e. putting aside the empirical estimates of individual objects’ weights). This analysis is described in S2 Supporting Information.

As described in the Introduction, Peters and colleagues (6) assume a prior over weight and density ratio that is comprised of three sub-priors, representing three different classes of weight-density relationships (here re-expressed as weight-volume relationships, see Figure 1C). Figure 7 compares the empirical biases in perceived log weight ratio (A) to the predicted biases from the best-fitting model parameters (B). Sub-priors (represented by dashed lines) are encoded as bivariate Gaussians (See Table E in S2 Supporting information for model parameters and model comparisons). Unfortunately, the model cannot account for the set of biases in perceived log weight ratio across all stimulus pairs (R^2^ = 0.413). It predicts a negative influence of log density ratio for some stimulus pairs; the predicted effect depends on where the stimuli lie relative to the three sub-priors. For example, in pairs of equal-volume (different weight, density) stimuli, log weight ratio is predicted to be underestimated as log density ratio, log weight ratio increase, whereas empirically it is overestimated.

**Fig 7.**
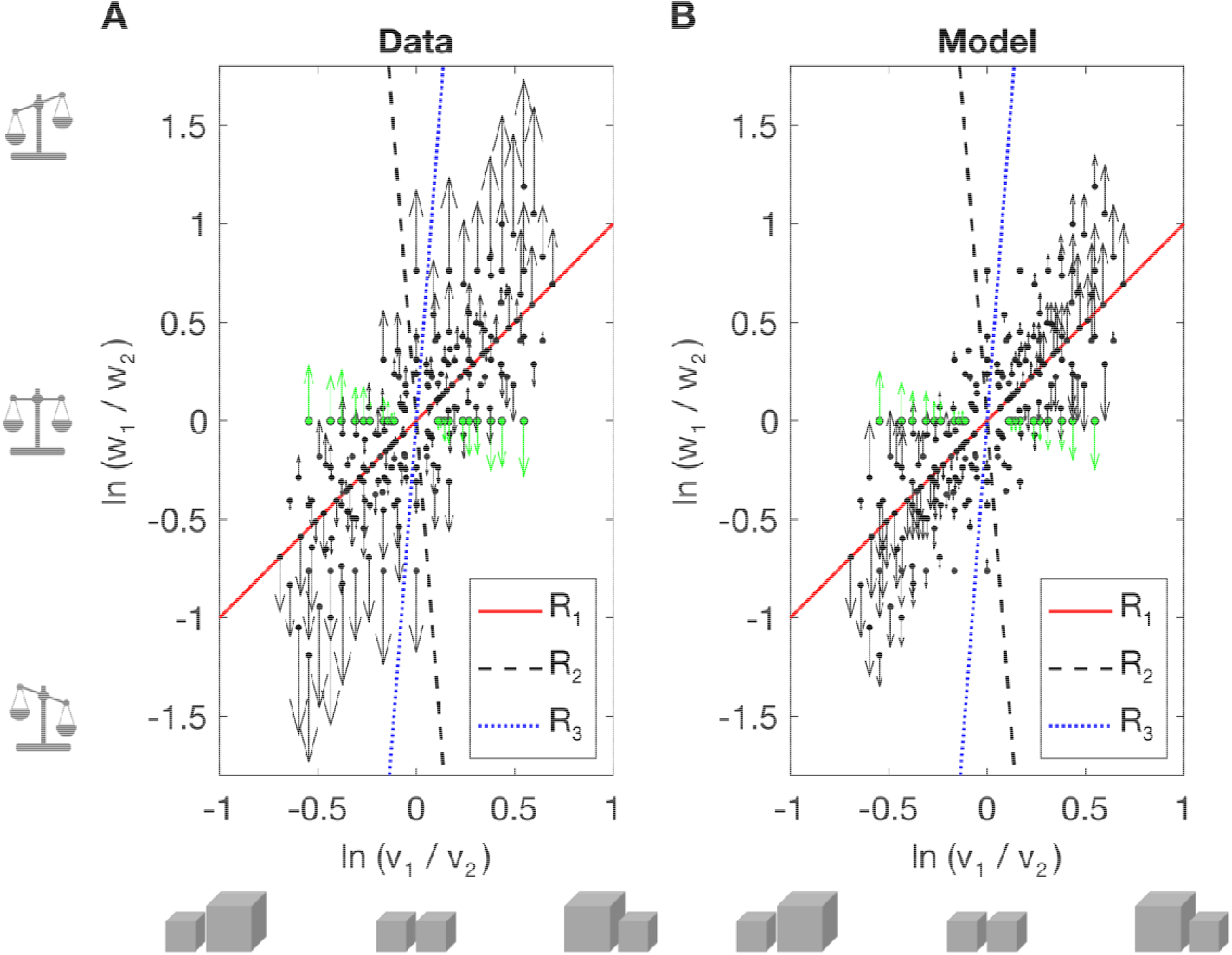
Peters and colleagues’ model of perceived weight. (A) Empirical biases in perceived log weight ratio. Dotted lines show the optimal three sub-priors from Peters and colleagues’ (6) model when fitted to our data. Black dots show the stimulus pairs; arrows show the magnitude and direction of the bias, with SWI conditions in green. (B) Predicted biases according to Peters and colleagues’ (6) model.

Finally, we evaluated Wolf and colleagues’ (11) model fit to our data. In this model, perceived weight is a weighted average of weight and density estimates, weighted according to their relative reliabilities. Estimates are modelled as power functions of the corresponding physical feature. As we didn’t manipulate the reliability of size information, the model reduces to:

The model provides a good account of perceived weight (R^2^ = .96). Figure F in S2 Supporting information shows the fit to our data, for comparison with Figure 3; for model parameters and model comparisons see Table F in S2 Supporting information). However, like Bays’ (28), it does not capture the contextual effects of the other object, including the observation the influence of density is strongly modulated by weight difference (as shown in Figure 4A).

We note that Wolf and colleagues (11) model perceived weight and density as power functions of their physical counterparts. However, additional exponent parameters for weight or density were not statistically supported by our data. Wolf and colleagues had a 5:1 ratio of heaviest to lightest stimuli, compared to our 3.3:1 ratio. It is possible that, given an increased range of stimulus weights / densities, our descriptive model would need to similarly include power functions of weight and / or density, in tandem with an intercept of 0.

## 6. Discussion

We investigated perceived weight in a set of stimuli which varied in weight and / or volume, and / or density in small steps. We report several novel findings: 1) The SWI occurs even at very small density (size) differences; 2) Across all stimuli, an object’s weight and density appear confounded / integrated to estimate weight; 3) Perceived weight is affected by the weight of the other object in a pair but is largely invariant to the other object’s density. 4) However, the influences of both objects’ densities on perceived weight are strongly modulated by weight difference. The negative, or ‘contrast’ effect of the other object’s density only appears for pairs of similar weight.

Our descriptive model captures the above effects, providing an excellent account of perceived weight (R^2^ = .98). It captures the SWI as well as biases in perceived weight across a range of variations in pairings of stimulus weight and density. Additionally, we evaluated a novel efficient-coding Bayesian model of perceived weight ratio, alongside previously proposed models of the SWI. Here we discuss our findings within the existing literature.

The magnitude of the SWI increased, as expected, with increasing density (size) difference (6,34,38,39). We extended this, by measuring the SWI with smaller density differences than those previously tested. We hypothesised that under small conflicts, the SWI might reverse (i.e., to show integration: smaller stimuli would be perceived as lighter). However, the SWI was clearly present for all equal-weight pairs, at odds with the idea that the SWI is driven by a (substantial) conflict between expected and perceived weight.

Our results are inconsistent with Peters and colleagues’ (6) Bayesian mixture model. Also, at a conceptual level, the subdivision of the continuous world of size and volume/density into three relatively narrow categories is difficult to reconcile with environmental statistics. The best fitting parameters for our data assign a prior probability of near 0 to equal-density objects (R_1_, see Figure 7, and Table E in S2 Supporting information). The other two sub-priors encapsulate the expectation that smaller objects are either much heavier and denser (R_2_) or much lighter and less dense than larger ones (R_3_), both at odds with reported natural statistics for everyday liftable objects (5).

The efficient-coding Bayesian ratio model provides a theoretically appealing account of the SWI, predicting a repulsion away from a plausible equal-density prior. However, neither our efficient-coding Bayesian ratio model, nor that recently proposed by Bays (28) satisfactorily account for our pattern of data. Whilst our efficient-coding model provides a reasonable fit to the empirical biases in perceived log weight ratio, it fails to account for the observed pattern of response variance, and does not explain biases in perceived weight for individual objects. As noted above, Bays’ model (28) does not account for contextual bias effects found in our data (i.e., the influence of the other object’s weight and, in certain conditions, its density), and neither is it consistent with empirical patterns of response variance.

So, is the SWI Bayesian after all? We don’t think so; the SWI does not appear to be caused by repulsion from an equal-density prior. It is reasonable to assume that the perceptual system does expect same-material objects to be same-density. Indeed, the sensorimotor system may rely on an object’s size, together with density expectations for classes of objects / materials to infer the weight of a stimulus prior to lifting (14). Such expectations presumably drive the initial (erroneous) forces used when lifting SWI objects (4). However, the illusion persists after forces quickly adapt to the true weights (4) and the SWI is just as large when same-density cues are reduced, i.e., when the colour or surface material of the two stimuli differs (54,70). Thus, while we may employ an equal-density prior, this does not seem to underlie the perceptual SWI.

In general, perceived weight was modulated by density; denser objects are perceived as heavier than less dense objects, consistent with previous literature (11). In line with Wolf and colleagues (10,11) weight appears to be routinely confounded / integrated with density information. This is consistent with previous evidence that the SWI still occurs when the smaller objects is slightly lighter (but still denser) than the larger object (18,46). However, our data reveal that the effect of density is substantially increased when equal-weight objects are encountered, as in the typical size-weight illusion paradigm. Thus, like Bays’ (28) model, Wolf and colleagues’ (11) model fails to capture the contextual effects of the other object. Rather, these models predict biases in perceived weight that occur in single-object judgements (see, e.g., (71)).

Our data and model reveal the role of contextual effects in the classic SWI (where weight is typically held constant) and highlight the error in generalising from these data to weight perception in general. Our model falls short of providing a complete model of weight perception, because we focussed on pairs of objects. Further work might determine how the model could be expanded to capture weight perception in a wider variety of contexts. For single objects presented in sequence, for example, one could replace the other object’s properties with a weighted combination of previously encountered stimuli.

Wolf and colleagues (11) also showed that when the reliability of (visual or haptic) size information is reduced, density has less effect on the final heaviness estimate. In the limiting case, when size information is completely absent (objects lifted via strings / a handle, and no vision), the illusion disappears (11,39,72). Our model of weight perception does not encapsulate the additional effect of varying the reliability of size estimation; this would be accommodated by reducing the coefficients associated with density to reflect the availability of visual-haptic size cues (*β*_2_, *β*_4_, *β*_5_, Equation 1).

## 7. Conclusions

Our study situates the SWI in the broader context of weight perception under varied conditions (combinations of volume, weight, and density) allowing us to evaluate different models of weight perception, including our own.

Our explorations reveal that, whilst the SWI magnitude increases with size / density difference, it occurs even for small density differences. Second, we showed that typical SWI experiments have inflated the apparent role of density on weight perception more generally. The effect of density on perceived weight is substantially smaller when two objects of different weights are encountered. Our simple model provides an excellent account of perceived weight under various combinations of size, weight, and density. However, we still don’t know why weight is estimated in this way. Why does density affect weight perception, and why is this effect magnified when weight differences are small?

We explored different avenues to reconcile the SWI within the Bayesian framework, given the appeal of finding a model that provides a more explanatory account of weight perception in the SWI and beyond. However, none of the Bayesian models we tested, including efficient-coding models that do predict biases away from a prior, could successfully account for our data.

## Acknowledgments

We thank Megan A K Peters for helpful discussions about modelling, and Benjamin K Graf for assistance with deriving the equations for predicted response variance (see S2 Supporting Information).

## Supporting information captions

**S1 Supporting Information. Control experiment**

**S2 Supporting Information. Other models**

