## Supplementary material for "The size-weight illusion and beyond: a new model of perceived weight": S1 Supporting Information

### S1 Supporting Information. Control experiment

In our main experiment, we expected that observers' perception of the weight difference between objects would be consistent with their weight estimates of the individual objects' weights. For example, if a participant judged one object to weigh 10 units and the other to weigh 15 units, we inferred that the perceived difference between the two would be 5 units. In other words, judgements within each stimulus pair would obey transitivity. However, failures in transitivity do occur in human perception. To test for transitivity within object pairs, we ran a brief control experiment. We asked 8 naïve participants who did not take part in the original experiment to report all three values (the two individual weights, and their difference) for a sample of stimulus pairs (N = 21) from our original experiment. Aside from these details, the methods were the same as in the main experiment. The experiment was approved by the University of Southampton Psychology Ethics Committee (No 101430); all participants gave written prior informed consent.

The stimulus set included the reference stimulus and two further stimuli from each stimulus subset, shown in Figure A (panel A), for comparison with Figure 2B in the main manuscript. All possible object pairings were presented twice each (with the left-right position of the objects switched across repetitions).

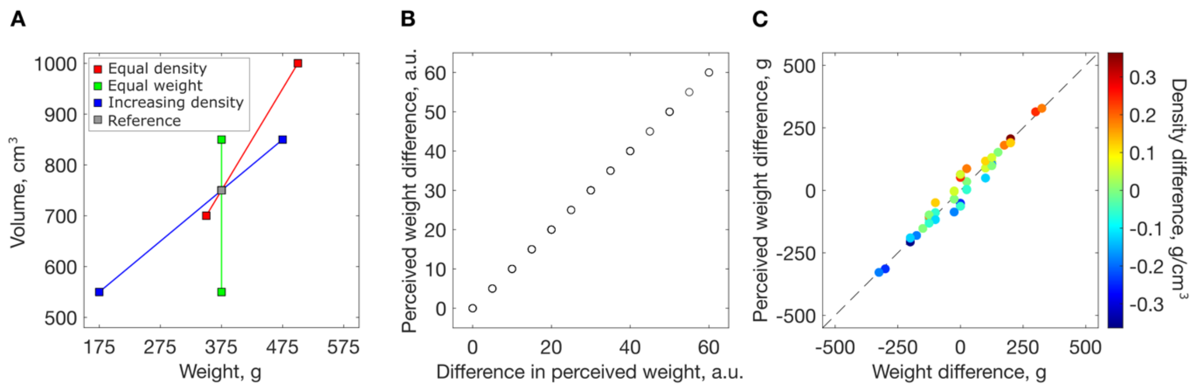


**Figure A. Control experiment.** (A) The stimulus set used in the control experiment. (B) The relationship between (i) the perceived weight difference between objects within a pair and (ii) the difference between individual estimates of the two objects' weights; a.u. = arbitrary units. (C) The perceived weight difference between objects as a function of their true weight difference (x-axis) and their density difference (indicated by colour).

On each trial (N = 42), participants reported the perceived weight difference between the two objects in the pair, in addition to the perceived weight of each object, as in the original experiment. Both were reported in the same arbitrary units. As in the main experiment, the reference was presented in isolation before the trials, and again after every 20 trials (The reference object also featured in some of the object pairs).

Figure A (panel B) plots the perceived difference in the weight of the two objects (explicitly reported) against the difference in the perceived weights (i.e., the difference between the separately reported magnitudes of the two objects' weights), for each trial / participant (336 datapoints). It is clear that the two are equivalent. This was true whether observers were first asked to report the perceived difference, and then asked for the individual weights, or vice versa.

Data from the control experiment are summarised in Figure A (panel C), which shows the perceived weight difference (or difference between the perceived weights) on the y-axis as a function of the true weight difference between the objects (x-axis) and their density difference (indicated by the colour). This can be compared with Figure 4A in the main manuscript. These results confirm that participants' responses in the two tasks are mutually consistent. In other words, we can confirm that individual weight estimates in our main experiment implicitly capture the perceived weight differences between objects pairs, as described by Equation 2 in the main text.

It is possible that participants mis-reported the ‘perceived difference’ to be consistent with their estimates of individual weights, even when those were yet to be made. However, the more parsimonious explanation is that the two types of perceptual judgements obey internal consistency. Indeed, a few participants spontaneously reported estimating individual weights first, in order to infer the difference, as they found direct difference judgments challenging or impossible.
