## Supplementary material for "The size-weight illusion and beyond: a new model of perceived weight": S2 Supporting Information

### S2 Supporting Information. Other models

#### 1. Alternative candidate models

We report alternative candidate models for perceived weight ($\hat{w}_1$) as a function of the judged and other object's weight and density (Table A) / volume (Table B) in the full dataset. Models of different complexity were compared using leave-one-out cross validation over participants (see Table A).

| **Model N** | 1 | 2 | 3 | 4 | 5 | 6** | 7 |
|---|---|---|---|---|---|---|---|
| **N parameters** | 2 | 3 | 4 | 5 | 6 | 7 | 8 |
| **Intercept** | -80.42 | -149.57 | -201.33 | -186.53 | -197.74 | -181.34 | -182.91 |
| $\beta_1$ | 1.29 | 1.18 | 1.19 | 1.20 | 1.21 | 1.23 | 1.23 |
| $\beta_2$ | | 219.70 | 219.70 | 216.54 | 172.87 | 126.43 | 131.26 |
| $\beta_3$ | | | 0.13 | 0.15 | 0.13 | 0.13 | 0.13 |
| $\beta_4$ | | | | -43.87 | 12742 | 27814 | 24454 |
| $\beta_5$ | | | | | | -17919 | -14510 |
| **σ ($\sigma_1$)** | | | | | 92.67 | 76.44 | 64.16 |
| $\sigma_2$ | | | | | | | 56.40 |
| **SSR** | 161917 | 113295 | 80075 | 78145 | 61372 | 56105 | 55592 |
| **$R^2$:**<br>**Est. $w_1$**<br>**Est. log w ratio**<br>**Bias in log w ratio** | <br>.9541 | <br>.9679<br>.9836<br>.8469 | <br>.9773 | <br>.9778 | <br>.9826 | <br>.9841<br>.9905<br>.9117 | <br>.9842 |
| **XVal MSE:**<br>**Est. $w_1$**<br>**Est. log w ratio** | <br>$1.088\times10^{6}$ | <br>$1.043\times10^{6}$ | <br>$1.012\times10^{6}$ | <br>$1.010\times10^{6}$ | <br>$9.947\times10^{5}$ | <br>$9.866\times10^{5}$<br>20.6271 | <br>$9.906\times10^{5}$ |
| **F-ratio test *p***<br>**Models compared** | | < .001<br>2 vs 1 | < .001<br>3 vs 2 | < .001<br>4 vs 3 | < .001<br>5 vs 3 | < .001<br>6 vs 5 | .71<br>7 vs 6 |

**Table A. Fitted parameters and model comparisons for alternative candidate models for perceived weight ($\hat{w}_1$).** The double asterisk indicates the best model. $w_1$, $w_2$ = judged and other object's weight; $d_1$, $d_2$ = judged and other object's density. The model with the lowest cross-validation error (XVal MSE, corresponding to the mean summed squared residuals, SSR over left-out participants) is generally accepted as the 'best' model, i.e. that which is expected to most accurately predict new participants' data. Nonetheless, F-ratio tests were performed to compare nested models (df = 105 - $N_{params}$).

Model 1: $C+\beta_1 w_1$

Model 2: $C+\beta_1 w_1+\beta_2 d_1$

Model 3: $C+\beta_1 w_1+\beta_2 d_1+\beta_3 w_2$

Model 4: $C+\beta_1 w_1+\beta_2 d_1+\beta_3 w_2+\beta_4 d_2$

Model 5: $C+\beta_1 w_1+\beta_2 d_1+\beta_3 w_2+\beta_4\, \mathrm{N}(|w_1-w_2|, 0, \sigma^2)$

Model 6: $C+\beta_1 w_1+\beta_2 d_1+\beta_3 w_2+(\beta_4+\beta_5)\mathrm{N}(|w_1-w_2|, 0, \sigma^2)$

Model 7: $C+\beta_1 w_1+\beta_2 d_1+\beta_3 w_2+\beta_4\, \mathrm{N}(|w_1-w_2|, 0, \sigma_1^2)+\beta_5\, \mathrm{N}(|w_1-w_2|, 0, \sigma_2^2)$

| **Model N** | 8 | 9 | 10 |
|---|---|---|---|
| **N parameters** | 3 | 4 | 5 |
| **C** | -146.22 | -109.09 | -109.09 |
| $\beta_1$ | 1.30 | 1.47 | 1.48 |
| $\beta_2$ | 0.08 | 0.08 | 0.05 |
| $\beta_3$ | | -0.13 | -0.13 |
| $\beta_4$ | | | 0.07 |
| **SSR** | 125152 | 93198 | 89943 |
| $R^2$ | .9645 | .9736 | .9745 |
| **XVal MSE** | $1.053\times10^6$ | $1.025\times10^6$ | $1.022\times10^6$ |
| **F-ratio test *p*** **Models compared** | < .001 8 vs 2 | < .001 9 vs 8 | < .001 10 vs 9 |

**Table B. Fitted parameters and model comparisons for alternative candidate models for perceived weight ($\hat{w}_1$).** $w_1$, $w_2$ = judged and other object's weight; $v_1$, $v_2$ = judged and other object's volume.
Model 8: $C+\beta_1 w_1+\beta_2 v_2$
Model 9: $C+\beta_1 w_1+\beta_2 v_2+\beta_3 v_1$
Model 10: $C+\beta_1 w_1+\beta_2 v_2+\beta_3 v_1+\beta_4 w_2$

Figure A shows weight estimates / difference in weight estimates for the SWI pairs as a function of volume / volume difference, for comparison with Figure 3C, D in the main text.

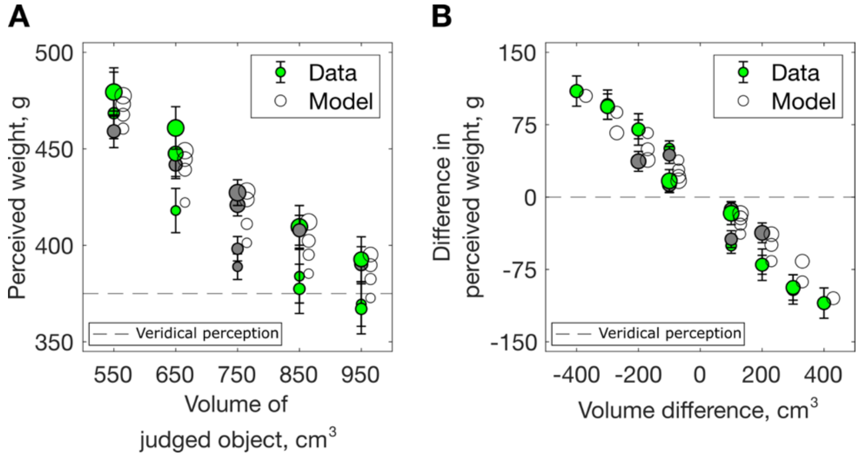


**Figure A. Results for SWI pairs re-expressed as a function of stimulus volume / volume difference.** (A) Perceived weight as a function of stimulus volume. (B) Difference in estimated weight as a function of volume difference. Data averaged across participants, error bars show ± 1 SEM. Open symbols show the model predictions (described in the main text, Equations 1 and 2). The dashed lines indicate veridical perception.

### Efficient-coding Bayesian ratio model

To fit the efficient-coding Bayesian ratio model, we did not model the full Bayesian inference process. Instead, we use Wei and Stocker (29, 30) equations for the bias (Equation 3 in the main text) and reliability (see below, Equation C). The model includes a prior over log weight (w) ratio and log volume (v) ratio, centred on [0, 0]. We compared models with priors of different complexity (either one bi-variate Gaussian, or a mixture of up to 4). Each component Gaussian ($G_1$–$G_4$) of the prior is defined by the three parameters of the covariance matrix: the variabilities of log w ratio and log v ratio and their correlation ($\sigma_{wr}$, $\sigma_{vr}$, $\rho$, respectively). For priors comprised of more than one Gaussian, we also fitted the weights (summing to 1) of each component (for $G_1$–$G_{N-1}$). Last, we fitted the bias coefficient (C) (see Equation 3). The bias coefficient is determined by factors including amount of internal and external noise, put in practice is fitted as free parameter. For each set of potential parameters, the 2D prior was computed. Then, using Equation 3, we found the expected bias for each log w ratio, log v ratio stimulus pair, from the values of the prior at (log w ratio - $\delta$, log v ratio) and (log w ratio + $\delta$, log v ratio). Optimal parameters (minimising the SSR biases in log weight ratio) were identified via gradient descent, using multiple different starting values. Fitted parameters and model comparisons are shown in Table C. Models were compared using leave-one-out cross validation over participants.

| **Model N** | 11 | 12 | 13* | 14 |
|---|---|---|---|---|
| **N parameters** | 4 | 8 | 12 | 16 |
| **$G_1$** | | | | |
| - $\sigma_{wr}$ | 2.532 | 0.504 | 3.354 | 0.395 |
| - $\sigma_{vr}$ | 0.761 | 0.262 | 1.056 | 0.270 |
| - ρ | 0.216 | 0.449 | -0.521 | 0.644 |
| - weight | - | 0.118 | 0.874 | 0.121 |
| **$G_2$** | | | | |
| - $\sigma_{wr}$ | | 5.835 | 0.280 | 1.430 |
| - $\sigma_{vr}$ | | 1.064 | 0.303 | 1.006 |
| - ρ | | -0.6 | 0.564 | 0.337 |
| - weight | | | 0.017 | 0.630 |
| **$G_3$** | | | | |
| - $\sigma_{wr}$ | | | 0.455 | 0.917 |
| - $\sigma_{vr}$ | | | 0.250 | 2.748 |
| - ρ | | | 0.576 | -0.267 |
| - weight | | | - | 0.223 |
| **$G_4$** | | | | |
| - $\sigma_{wr}$ | | | | 0.55 |
| - $\sigma_{vr}$ | | | | 2.748 |
| - ρ | | | | 0.964 |
| - weight | | | | - |
| **C** | $1.7173\times10^{10}$ | $2.0141\times10^{11}$ | $1.3665\times10^{11}$ | $1.2835\times10^{11}$ |
| **SSR** | 1.1698 | 0.9042 | 0.8386 | 0.8034 |
| **$R^2$** | | | | |
| **Est. log weight ratio** | .9836 | .9873 | .9883 | .9887 |
| **Bias in log w ratio** | .8472 | .8819 | .8905 | .8951 |
| **XVal MSE** | 21.0205 | 20.7945 | 20.7180 | 20.7378 |
| **F-ratio test *p*** | | < .001 | .002 | .55 |
| **Models compared** | | 12 vs 11 | 13 vs 12 | 14 vs 13 |

**Table C. Fitted parameters and model comparisons, given priors of different complexity, for the efficient-coding Bayesian ratio model.** The asterisk indicates the best model among those reported in the table.

Using the optimal fitted parameters, we generated predictions for biases in perceived weight of the two objects, $w_1$ and $w_2$ in each pair (see Figure B). Predictions were derived by splitting the fitted bias in log weight ratio between the two stimuli such that the perceived log weights of $w_1$ and $w_2$ are under- and over-estimated by the same percentage. It is clear that the observed (panels A, B) and predicted (panels C, D) biases follow very different patterns.

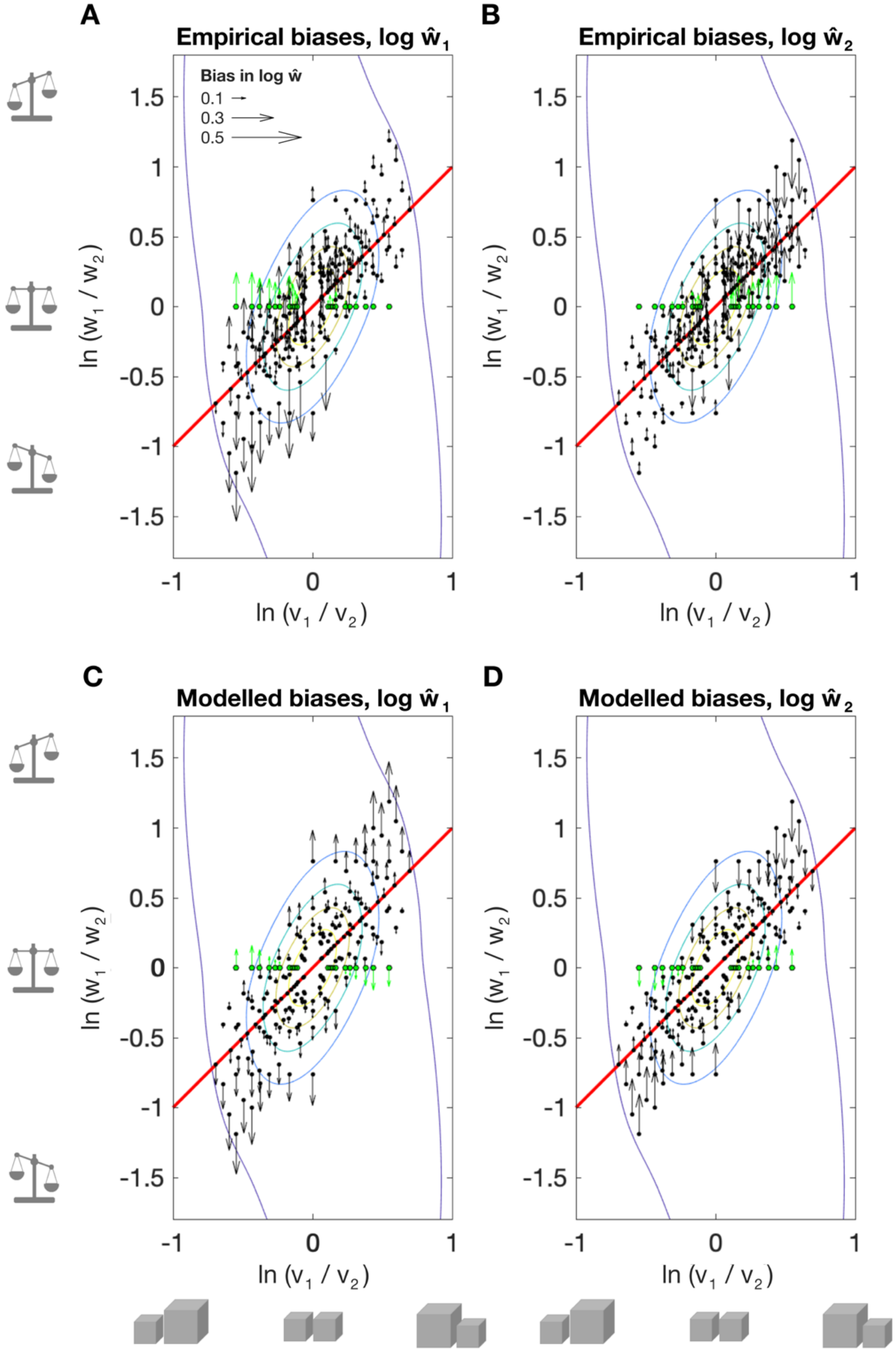


**Figure B. Efficient Bayesian ratio model, data and predictions for perceived log weight ($\log \hat{w}_1$) of the two individual objects in each pair.** The true log weight ratio and log volume ratio of the pair are given by black dots. The length of the arrow gives the magnitude of bias in perceived log weight (see legend in A) of $w_1$ (left plots) or $w_2$, (right plots) and its direction gives the sign (up = positive, down = negative). (A) Empirical biases in perceived log $w_1$ and (B) log $w_2$. We assume the prior to have equal and opposite effects on perceived weight of the two objects; these predictions are shown below (C, D).

From Wei & Stocker (30) the relationship between the prior and the perceptual bias is given by $b(\theta) = C_b(1/p(\theta)^2)'$ (where $\theta$ is the stimulus distribution) and the response variance $\hat{\sigma}^2(\theta)$ is related to perceptual bias as $b(\theta) \propto (\hat{\sigma}^2(\theta)/(1+b'(\theta))^2)'$. Substituting for $b(\theta)$ in these equations gives:

$$C_\sigma \left(\frac{1}{p(\theta)^2}\right)' = \left(\frac{\hat{\sigma}^2(\theta)}{\left(1+b'(\theta)\right)^2}\right)'. \qquad \text{(A)}$$

$$\frac{C_\sigma}{p(\theta)^2} = \frac{\hat{\sigma}^2(\theta)}{\left(1+b'(\theta)\right)^2} + C_0. \qquad \text{(B)}$$

Because $\hat{\sigma}^2 \rightarrow 0$ as $p(\theta) \rightarrow \infty$ we can set $C_0 = 0$, and so

$$\hat{\sigma}^2(\theta) = C_\sigma(1+b'(\theta))^2/p(\theta)^2. \qquad \text{(C)}$$

Figure C compares the empirical response variance, vs. the predictions (Equation C). Empirically, response variance is proportional to log weight ratio, but this is not predicted by the model. The prediction is that response variance will be primarily modulated by the log volume ratio.

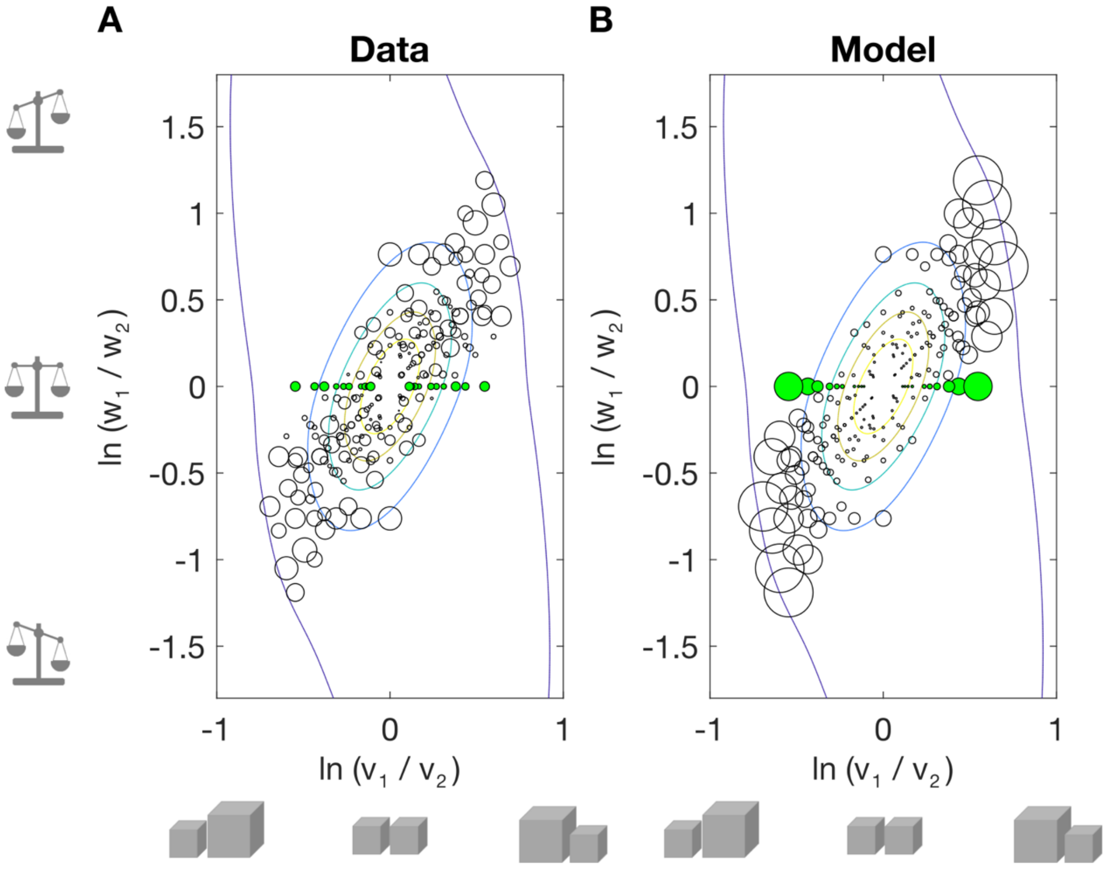


**Figure C. Variance for estimates of log weight ratio.** (A) Empirical variances, (B) variances predicted by our efficient-coding Bayesian model. The location of each circle gives the true log volume ratio and log weight ratio of the stimulus pair. SWI pairs are filled in green. The circle radius is proportional to the within-observer variance of the ratio of the two objects' estimated weights, averaged across observers. In the empirical data, response variance is modulated by the absolute log weight ratio (most reliable judgements occur when the object weights are similar). In the model predictions, response variance is primarily modulated by log *volume* ratio.

We also considered Bayesian models involving a 2D prior over *differences* in weight and volume or density (i.e. $w_1 - w_2$, $v_1 - v_2$, $d_1 - d_2$). A prior / model defined with regards to weight and volume differences is problematic: knowing the weight difference and volume difference of a stimulus pair leaves the density difference undetermined. This model thus (nonsensically) gives identical predictions of bias for a given weight difference, irrespective of whether the density difference is positive or negative. In principle, a prior over weight and *density* difference is more plausible, given that our descriptive model predicts perceived weight difference as a function of weight difference and density difference. In practice, however, this model provides a worse fit to the observed biases in perceived weight difference (mixture of 2 Gaussian priors, 8 parameters; $R^2$ = .7662, SSR = 88230) than our descriptive model (which reduces to 4 parameters when predicting perceived weight differences, see Equation 2 in the main text; $R^2$ = .9117, SSR = 1025.1). Moreover, the model predicts a completely different pattern of variance in estimated weight difference than the one observed ($R^2$ = .0001, p = .89).

**S4. Bays' efficient coding Bayesian model**

Bays' model (28) includes a prior over weight ($w_1$) and volume ($v_1$), for single objects. Our extension of the model uses a mixture of bi-variate Gaussians, rather than just one. As for our efficient coding Bayesian ratio model (see above), we compared models with priors of different complexity (1 to 4 Gaussians). Each Gaussian component of the prior ($G_1$–$G_4$), is defined by the mean ($\mu_w$, $\mu_v$) and variability ($\sigma_w$, $\sigma_v$,) of $w_1$ and $v_1$ and their correlation (ρ). For priors comprised of more than one Gaussian, we also fitted their respective weights ($G_1$–$G_{N-1}$). We also fitted the bias coefficient (C) (see Equation 3 in the main text). For each set of potential parameters, the prior was computed. The expected bias for each $w_1$ $v_1$, stimulus was then calculated using Equation 3, from the value of the prior at ($\log w_1 - \delta$, $\log v_1$) and ($\log w_1 + \delta$, $\log v_1$). Optimal parameters (minimising the SSR) were identified via gradient descent, using multiple different starting values. Table D provides fitted parameters and model comparisons.

| **Model number** | 15 | 16 | 17* | 18 |
|---|---|---|---|---|
| **N parameters** | 6 | 12 | 18 | 24 |
| **$G_1$** | | | | |
| - $\sigma_w$ | 5108.8 | 160.98 | 135.79 | 972.31 |
| - $\sigma_v$ | 671.41 | 719.82 | 436.49 | 161.14 |
| - ρ | 0.014 | 0.385 | 0.21 | 0.220 |
| - $\mu_w$ | 339.7 | 271.5 | 278.8 | 339 |
| - $\mu_v$ | 1262.2 | 62.9 | 56.3 | 995.4 |
| - weight | - | 0.049 | 0.016 | 0.186 |
| **$G_2$** | | | | |
| - $\sigma_w$ | | 1607.6 | 2153.2 | 55.56 |
| - $\sigma_v$ | | 481.21 | 259.98 | 182.21 |
| - ρ | | 0.49 | 0.47 | 0.319 |
| - $\mu_w$ | | 65.0 | 304.9 | 423.6 |
| - $\mu_v$ | | 658.6 | 680.4 | 1378.3 |
| - weight | | - | 0.223 | 0.015 |
| **$G_3$** | | | | |
| - $\sigma_w$ | | | 1605.7 | 424.49 |
| - $\sigma_v$ | | | 650.73 | 801.81 |
| - ρ | | | 0.18 | 0.75 |
| - $\mu_w$ | | | -9.8 | 513.81 |
| - $\mu_v$ | | | 289.6 | 383.43 |
| - weight | | | - | 0.144 |
| **$G_4$** | | | | |
| - $\sigma_w$ | | | | 656 |
| - $\sigma_v$ | | | | 682.76 |
| - ρ | | | | 0.56 |
| - $\mu_w$ | | | | 263.12 |
| - $\mu_v$ | | | | 1012.9 |
| - weight | | | | - |
| **C** | 1.503 | 0.031 | 0.072 | 0.032 |
| **SSR** | 96548.73 | 81136.29 | 80069.23 | 79719.90 |
| **$R^2$** | | | | |
| **Est. $w_1$** | .9726 | .9770 | .9773 | .9774 |
| **Bias in $w_1$** | .7112 | .7574 | .7605 | .7615 |
| **XVal MSE** | $1.0281\times10^{6}$ | $1.0139\times10^{6}$ | $1.0135\times10^{6}$ | $1.0137\times10^{6}$ |
| **F-ratio test *p*** | | < .001 | > .99 | > .99 |
| **Models compared** | | 6 vs 15 | 17 vs 16 | 18 vs 17 |

**Table D. Model comparisons, given priors of different complexity (1-, 2-, 3, 4-Gaussian priors), for Bays' efficient-coding Bayesian model fitted to our data.** Models were compared using leave-one-out cross validation over participants. The asterisk indicates the best model (according to the XVal MSE) among those reported in the table.

**Figure D** compares the empirical biases vs. the predictions for the best model fitted to our data (see Table D).

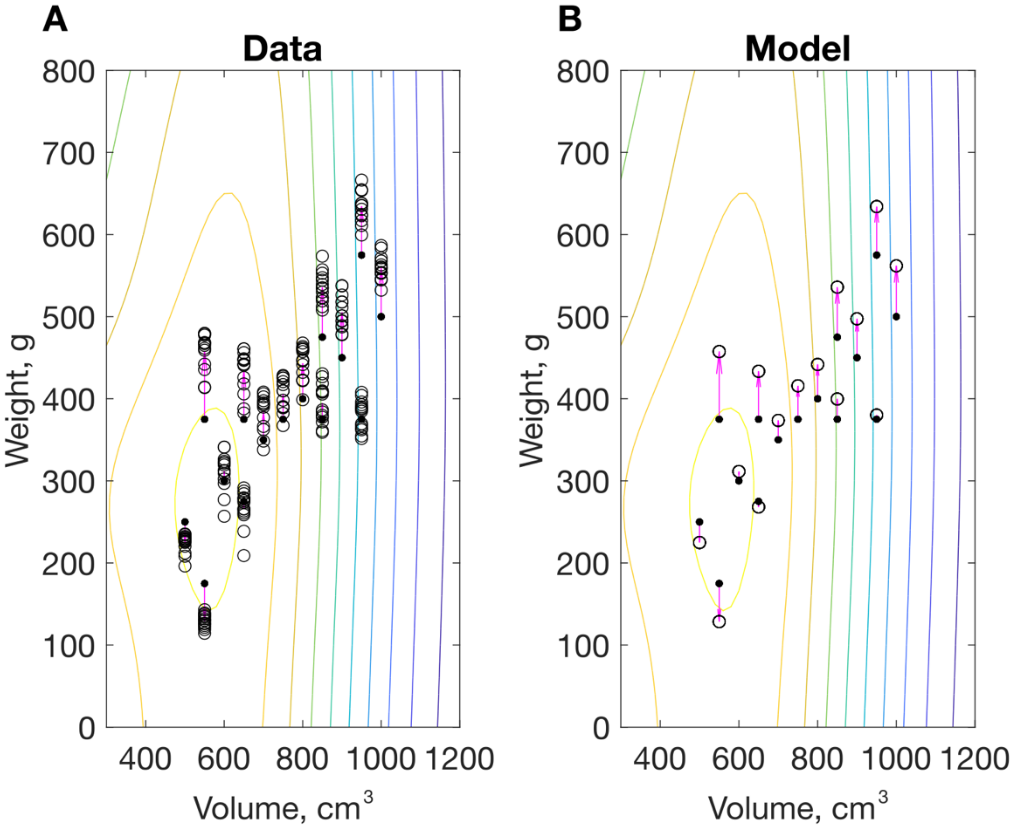


**Figure D. Biases in perceived weight ($\hat{w}_1$): empirical (A) and fits (B) for Bays' extended model fitted to our data.** Contours show the prior. Black dots show the stimuli; black circles (connected by magenta arrows) show observed (A) and predicted (B) estimates of $w_1$ according to the model.

Bays' model does not predict the observed pattern of response variance ($R^2 = 0.041$ $p < .01$, see Figure E). Rather, response variance is significantly (although weakly) correlated with stimulus weight (0.087, $p < .001$). In fact, Bays (28) infers the bias and variability of estimates of individual stimuli from judgements of weight ratios (data from (6)). For this, he relies on an assumption that is not supported by our data: that the weight estimates of the two constituent objects are independent.

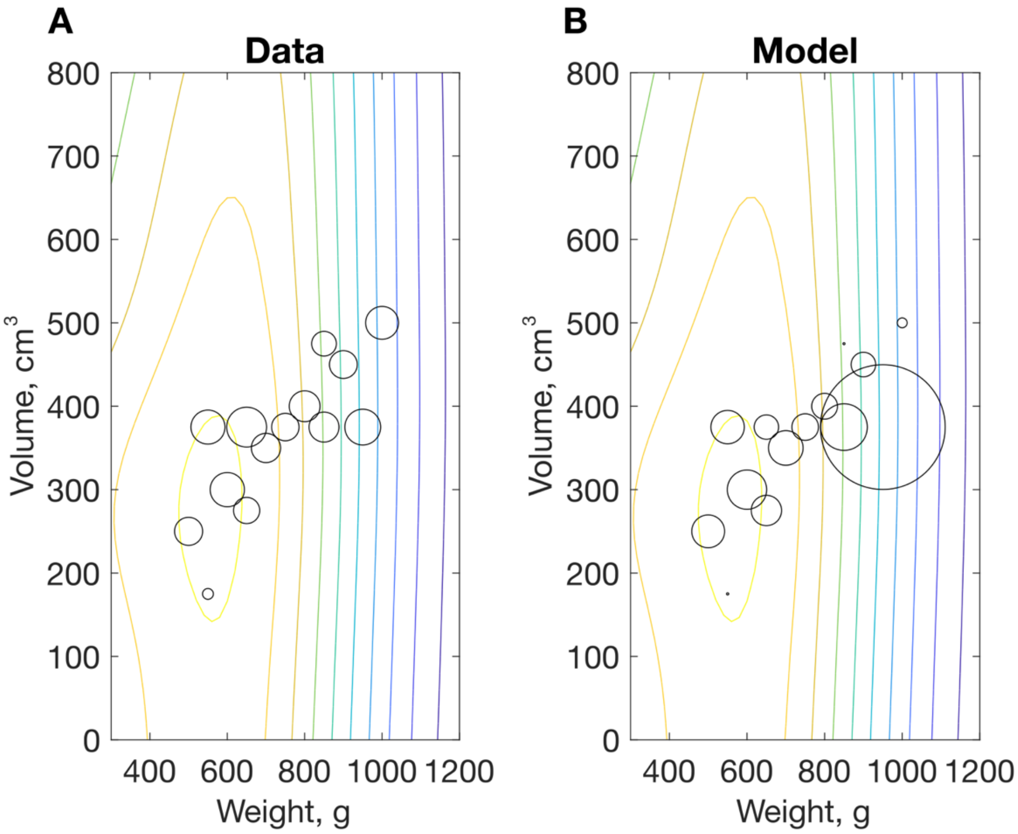


**Figure E. Response variance; empirical (A) and fits (B) for Bays' extended model fitted to our data (see Equation C).** The location of each circle gives the true weight and volume of the stimulus. The circle radius is proportional to the within-observer variance of the object's estimated weight, averaged across observers.

As noted above, Bays' model predicts biases in the estimated weights of single objects, given a prior over the weight and volume of single objects. The above analysis, therefore, assesses how well the model accounts for our participants' perception of individual objects. However, a proponent of Bays' model might suggest fitting it directly to the log weight ratio of estimate pairs, ignoring the empirical estimates of the single objects in each pair, and instead assuming that they are independent of each other (although this assumption is not supported by the data). This mirrors the approach in Bays (28) who fits Peters and colleagues' (6) dataset, in which perceived weights of individual objects are not available. Directly fitting the biases in log weight ratios (i.e. minimising the squared error between the predicted and observed log weight ratios) provides a very good fit to those ratios ($R^2$ = .919, SSR = 0.622; cross validation supports a model with 4 Gaussian components to the prior). (The fit is better, in fact, than the simpler efficient coding Bayesian ratio model ($R^2$ = .895); in Bays' model, both the shape of the prior and its location are free parameters, whereas in the efficient coding Bayesian ratio model the prior is centred on $\log(w_1/w_2)$, $\log(v_1/v_2)$ = [0, 0], for theoretical reasons). Importantly, however, Bays' model provides a very poor account of the variance in empirical log weight ratios ($R^2$ = .000; compare to $R^2$ = .444 for the Bayesian ratio model). Moreover, the model's predictions of biases in *individual objects'* perceived weights are poor ($R^2$ = .976, with huge residuals: SSR = $5.6115 \times 10^6$). The fit for individual objects is inevitably worse than when Bays' model is fit directly to those data, as above. Unfortunately, therefore, the Bays model remains an implausible account of perceived weight.

### S5. Peters and colleagues' Bayesian model

We fitted Peters and colleagues' (6) Bayesian model to either (i) our equal-density pairs only, or (ii) our full dataset data (log perceived weight ratio judgements), with fitted parameters shown in Table E. The model includes three sub-priors (see Figure 1B in the main text) over log volume ratio and log density ratio (although plotted over log weight and log volume ratio in Figures 1C and 7 in the main text). Each sub-prior is defined by a bi-variate Gaussian, centred on [0, 0], and with variability parameters $\sigma_v$, $\sigma_d$, and correlation ρ. The equal-density prior is essentially a delta function, defined by a very small $\sigma_d$ ($0.1 \times 10^{-10}$) and a nominal ρ of 0. Sub-priors 2 and 3 ($R_2$, $R_3$) share common variability parameters ($\sigma_v$ and $\sigma_d$), and were constrained to have equal and opposite ρ. The three priors have a priori probabilities that sum to 1, characterised by two weights ($weight_1$ and $weight_2$). Optimal parameters (minimising the SSR in log weight ratio) were identified via gradient descent, with many different starting parameters.

| | Equal-weight pairs | Full dataset |
|---|---|---|
| **Model number** | 19 | 20 |
| **N parameters** | 7 | 7 |
| **$\sigma_v$** | 0.67 | 0.07 |
| **$\sigma_d$** | 1.08 | 0.63 |
| **ρ** | 0.9993 | 0.74 |
| **$weight_1$** | 0.0003 | $3.4694 \times 10^{-21}$ |
| **$weight_2$** | 0.85 | 0.44 |
| **$\sigma_{est\ w}$** | 0.34 | 0.15 |
| **$\sigma_{est\ v}$** | 0.03 | 0.01 |
| **SSR** | 0.05 | 4.54 |
| **$R^2$** | | |
| **Est. log weight ratio** | .9498 | .9447 |
| **Bias in log w ratio** | .9498 | .4127 |
| **XVal MSE** | - | 23.4209 |

**Table E. Fitted parameters and model comparisons for Peters and colleagues competitive prior Bayesian model fitted to our data**. Equal-weight pairs only or full dataset.

The model provides a good fit to our data when only equal-weight (SWI) pairs are included ($R^2$ = .949); whilst we report larger SWIs than Peters and colleagues (our observers grasped the objects directly rather than via handles; see (11)), this can be accommodated in the model by increasing the slope of sub-prior $R_2$ (smaller is denser and heavier). However, the fit to the full set of biases is poor ($R^2$ .41, compare with model 6, Table A).

**S6. Wolf and colleagues' model**

Table F shows fitted parameters and model comparison statistics for Wolf and colleagues' (11) model (see Equation 4 in the main text) fitted to weight estimates ($\hat{w}_1$) in our dataset. Optimal parameters (minimising the SSR in estimated $w_1$) were identified via gradient descent. Figure F shows weight estimates for within-subset pairs alongside model fits (for comparison with Figure 3 in the main text).

| **Model number** | 21 |
|---|---|
| **N parameters** | 4 |
| **$\beta_1$** | 0.32 |
| **x** | 1.19 |
| **$\beta_2$** | 297.79 |
| **y** | 3.45 |
| **SSR** | 136115 |
| **$R^2$** | .9620 |
| **XVal MSE** | $1.0651 \times 10^6$ |

**Table F. Fitted parameters and model comparison statistics for Wolf and colleagues' model.** See Equation 4 in the main text.

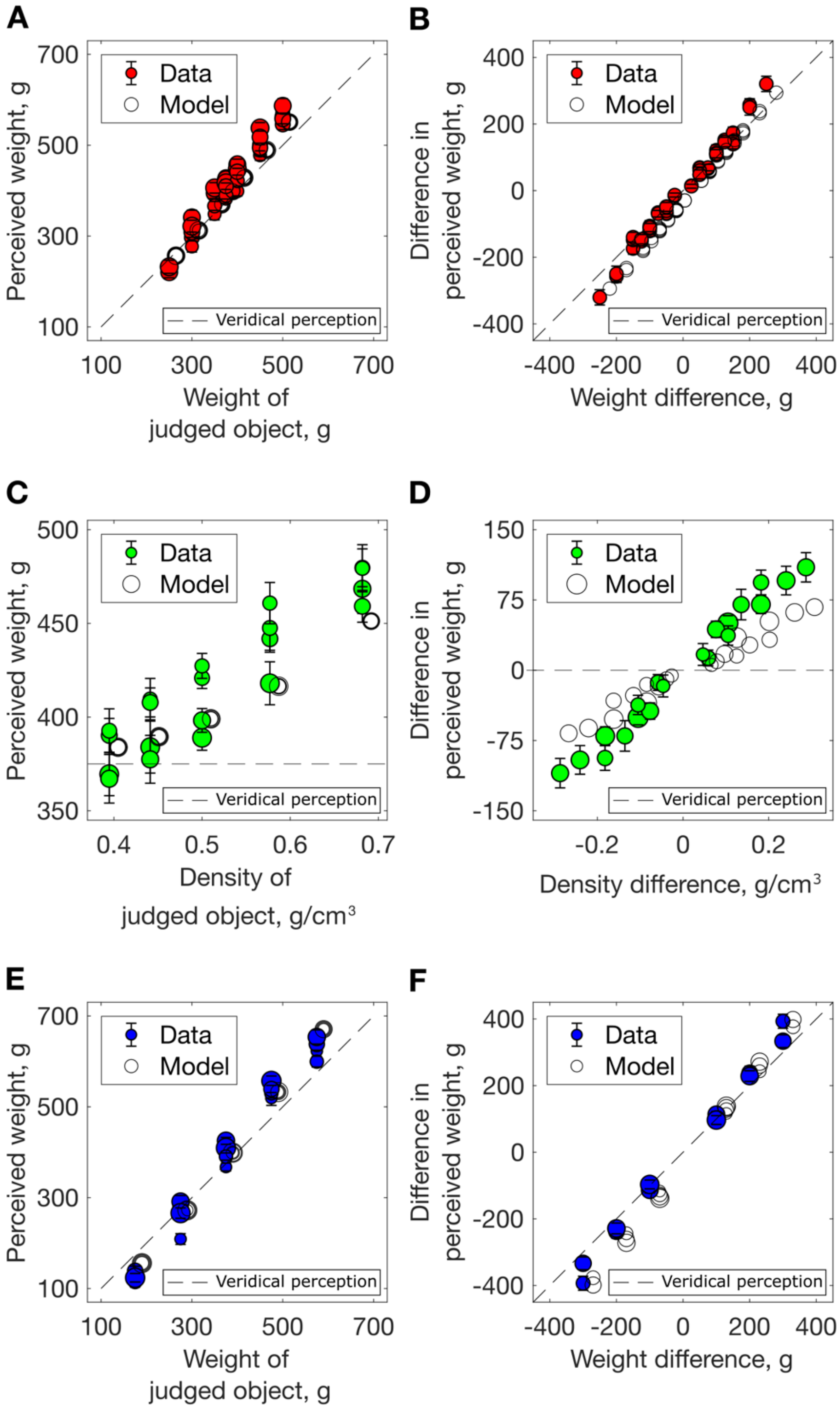


**Figure F. Data for within-subset pairs from our experiment with predictions from Wolf and colleagues' model**. Perceived weight (A, C, E) difference in perceived weight (B, D, F) for within-subset stimulus pairs, averaged across participants, error bars show ± 1 SEM. Solid lines / open symbols show the model predictions (see Equation 4 in the main text). The dashed lines indicate veridical perception. (A–B) Equal density pairs: (A) Weight estimates as a function of stimulus weight; symbol size indicates the weight of the other stimulus in the pair. (B) Data re-expressed as difference in perceived weight, as a function of weight difference. (C, D) Equal weight (SWI) pairs: (C) Weight estimates as a function of stimulus density; symbol size indicates the density of the other stimulus in the pair. (D) Data re-expressed as difference in perceived weight as a function of density difference; symbol size indicates the average object density. (E, F): Increasing density pairs: (E) Symbol size indicates the other stimulus' weight; (F) Data re-expressed as difference in perceived weight, as a function of weight difference; symbol size indicates the average object density.
